# Integrating barcoded neuroanatomy with spatial transcriptional profiling reveals cadherin correlates of projections shared across the cortex

**DOI:** 10.1101/2020.08.25.266460

**Authors:** Yu-Chi Sun, Xiaoyin Chen, Stephan Fischer, Shaina Lu, Jesse Gillis, Anthony M. Zador

**Affiliations:** Cold Spring Harbor Laboratory, Cold Spring Harbor, NY 11724, USA

## Abstract

Functional circuits consist of neurons with diverse axonal projections and gene expression. Understanding the molecular signature of projections requires high-throughput interrogation of both gene expression and projections to multiple targets in the same cells at cellular resolution, which is difficult to achieve using current technology. Here, we introduce BARseq2, a technique that simultaneously maps projections and detects multiplexed gene expression by *in situ* sequencing. We determined the expression of cadherins and cell-type markers in 29,933 cells, and the projections of 3,164 cells in both the mouse motor cortex and auditory cortex. Associating gene expression and projections in 1,349 neurons revealed shared cadherin signatures of homologous projections across the two cortical areas. These cadherins were enriched across multiple branches of the transcriptomic taxonomy. By correlating multi-gene expression and projections to many targets in single neurons with high throughput, BARseq2 provides a path to uncovering the molecular logic underlying neuronal circuits.

## Introduction

Neural circuits are comprised of neurons diverse in many properties, such as morphology (Lin et al., 2018; Winnubst et al., 2019), gene expression (Hodge et al., 2019; Saunders et al., 2018; Tasic et al., 2016; Tasic et al., 2018; Zeisel et al., 2018; Zeisel et al., 2015), and projections (Chen et al., 2019; Han et al., 2018; Harris et al., 2019; Wang et al., 2019). Although recent technological advances have made it possible to characterize the diversity in individual neuronal properties, associating multiple properties in single neurons with high throughput remains difficult to achieve. Investigating the relationship between multiple neuronal properties is essential for understanding the complex organization of neural circuits.

Of particular interest is the relationship between endogenous gene expression and long-range projections in the cortex. Cortical neurons have diverse patterns of long-range projections (Chen et al., 2019; Han et al., 2018; Huang et al., 2020; Oh et al., 2014; Wang et al., 2019) and diverse patterns of gene expression (Hodge et al., 2019; Scala et al., 2020; Tasic et al., 2016; Tasic et al., 2018; Yao et al., 2020; Zeisel et al., 2018; Zeisel et al., 2015; Zhang et al., 2020). The diversity in gene expression can be described by clustering neurons into transcriptomic types, but transcriptomic types have limited power in explaining the diversity of cortical projections beyond the major classes of projection neurons [(Chen et al., 2019; Klingler et al., 2018; Tasic et al., 2018; Zhang et al., 2020), but also see (Economo et al., 2018; Kim et al., 2019)]. The lack of clear correspondence between transcriptomic types and projections in the cortex raises the need to explore and identify other gene correlates of projections, potentially independent of transcriptomic types.

One class of candidate genes that might explain the diversity of projections is the cadherin superfamily, which we will refer to generally as “cadherins.” Cadherins are differentially expressed across cortical layers (Hayano et al., 2014; Krishna et al., 2009; Krishna et al., 2011; Redies, 1997) and cardinal inhibitory cell types defined by transcriptomic and phenotypic characteristics (Paul et al., 2017). Functionally, cadherins are known to specify and maintain neuronal connectivity (Basu et al., 2015; Duan et al., 2014; Duan et al., 2018; Hayano et al., 2014; Paul et al., 2017; Redies, 1997). Based on these studies, we hypothesize that the expression of specific cadherin superfamily members is correlated with specific patterns of projections (Fig. 1A). To test this hypothesis, we require a high-throughput technique that allows simultaneous multiplexed gene detection with projection mapping to multiple target areas at single-neuron resolution. Although advances in spatial transcriptomics have allowed high throughput and multiplexing capacity, achieving both multiplexed gene detection and high-throughput projection mapping in the same neurons remains difficult.

**Figure 1.**
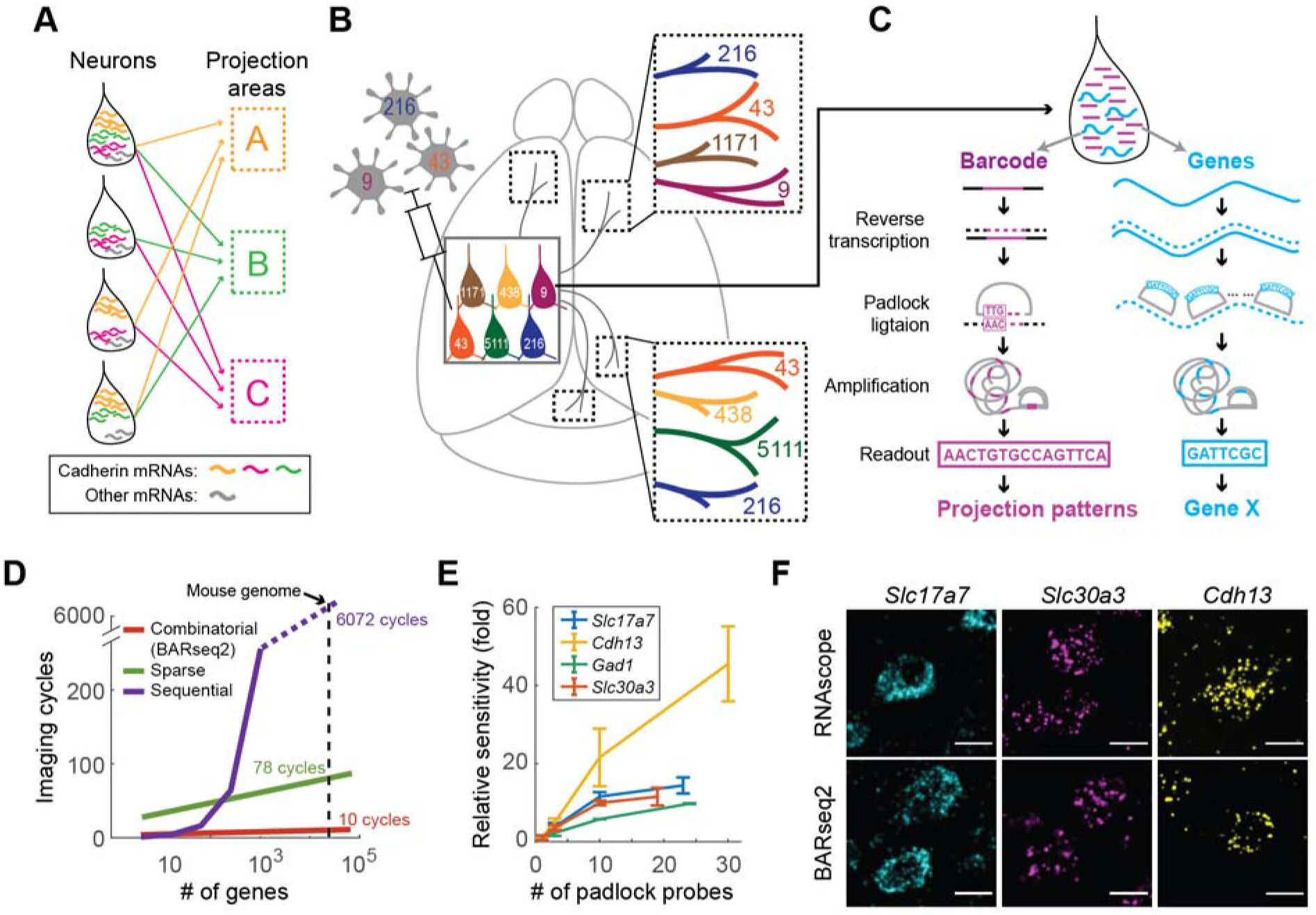
*In situ* sequencing of endogenous mRNAs using BARseq2. (A) Working hypothesis: the expression of cadherin mRNAs correlates with projections to diverse brain areas. (B)(C) BARseq2 correlates projections and gene expression at cellular resolution. In BARseq2, neurons are barcoded with random RNA sequences to allow projection mapping, and genes are also sequenced in the same barcoded neurons. RNA barcodes and genes are amplified and read out using different strategies (C). (D) Theoretical imaging cycles using combinatorial coding (BARseq2), 4-channel sequential coding, or 4-channel sparse coding as used by Eng et al. (2019). Imaging cycles assumed 3 additional cycles for BARseq2, 1 additional round for sparse coding, and no extra cycle for sequential coding for error correction. (E) Mean ± standard error of the relative sensitivity of BARseq2 in detecting the indicated genes using different numbers of padlock probes per gene. The sensitivity is normalized to that using one probe per gene. n = 2 slices for each gene. (F) Representative images of BARseq2 (*bottom*) detection of the indicated genes using the maximum number of probes shown in (E) compared to RNAscope (*top*). Scale bars = 10 μm.

To achieve high-throughput projection mapping, we recently introduced BARseq (<u>B</u>arcoded <u>A</u>natomy <u>R</u>esolved by <u>s</u>equencing), a projection mapping technique based on *in situ* sequencing of RNA barcodes (Chen et al., 2019). In BARseq, each neuron is labeled with a unique virally encoded RNA barcode that is replicated in the somas and transported to the axon terminals. The barcodes at the axon terminals located at various target areas are sequenced and matched to somatic barcodes, which are sequenced *in situ*, in order to determine the projection patterns of each labeled neuron. This sequencing-based projection mapping strategy, shared by both BARseq and a related technique, MAPseq (Kebschull et al., 2016), has been repeatedly validated using conventional neuroanatomical methods, and the contribution of potential artifacts—arising from sensitivity, double labeling of neurons, degenerate barcodes, fibers of passage, etc.—has been assessed and quantified in a variety of different systems using multiple techniques (Chen et al., 2019; Han et al., 2018; Huang et al., 2020; Kebschull et al., 2016).

Because BARseq preserves the location of somata with high spatial resolution, in principle it provides a platform to combine projection mapping with other neuronal properties also interrogated *in situ*, including gene expression. We have previously shown (Chen et al., 2019) that BARseq can be combined with fluorescent *in situ* hybridization (FISH) and *Cre*-labeling to uncover projections across neuronal subtypes defined by gene expression. However, these approaches can only interrogate one or a few genes at a time, which would be insufficient for associating a superfamily of cell adhesion molecules to diverse cortical projections.

To identify cadherin superfamily members that correlate with projections in the cortex, we aim to develop a technique to simultaneously map projections to multiple brain areas and detect the expression of dozens of genes in hundreds to thousands of neurons from a cortical area with high throughput, high spatial resolution, and cellular resolution. To achieve this goal, we combine the high throughput and multiplexed projection mapping capability of BARseq with state-of-the-art spatial transcriptomic techniques with high imaging throughput and multiplexing capacity (Ke et al., 2013; Qian et al., 2020). This second-generation BARseq (BARseq2) greatly improves the ability to correlate the expression of many genes to projections to many targets in the same neurons.

As a proof-of-principle, we first demonstrate multiplexed gene detection using BARseq2 by mapping the spatial pattern of up to 65 cadherins and cell-type markers in 29,933 cells. We then correlate the expression of 20 cadherins to projections to up to 35 target areas in 1,349 neurons in mouse motor and auditory cortex. Our study reveals novel sets of cadherins that correlated with homologous projections in both cortical areas. BARseq2 thus bridges transcriptomic signatures obtained through spatial transcriptional profiling with sequencing-based projection mapping to illuminate the molecular logic of long-range projections.

## Results

To investigate how cadherin expression relates to diverse projections, we developed BARseq2 to combine high-throughput projection mapping with multiplexed detection of gene expression using *in situ* sequencing (Fig. 1B, C). BARseq2 is based on BARseq, which achieves high-throughput projection mapping by *in situ* sequencing of RNA barcodes (Chen et al., 2019). In BARseq (Fig. 1C, *left*), RNA barcodes are reverse-transcribed and hybridized with a padlock probe that is complementary to the region flanking the barcode region. The barcode sequence is then copied into the padlock probe by a DNA polymerase, effectively “gap-filling” the padlock, and is subsequently ligated. After rolling circle amplification of the circularized padlock, the amplified RNA barcodes are then sequenced *in situ* using Illumina sequencing chemistry and matched to barcodes at target areas to identify projections (Fig. 1B). Projection patterns observed using RNA barcoding are consistent with those obtained using conventional neuroanatomical techniques in multiple circuits, including the locus coeruleus (Kebschull et al., 2016), auditory cortex (Chen et al., 2019), visual cortex (Han et al., 2018), and interregional connectivity across the whole cortex (Huang et al., 2020). Possible technical concerns, including distinguishing fibers of passage from axonal termini, sensitivity, double labeling of neurons, and degenerate barcodes, have previously been addressed and will not be discussed in detail again here. In particular, BARseq in both auditory cortex (Chen et al., 2019) and motor cortex (Chen et al., unpublished observations) produced single-cell projection patterns consistent with conventional retrograde tracing and single-cell tracing, while achieving throughput at least two to three orders of magnitude higher than the current state-of-the-art single-cell tracing techniques. For example, we were able to map up to 5,000 neurons per person-week (Chen et al., unpublished observations), which allowed us to uncover organizational principles of projections that would have been difficult to discover using smaller datasets. Combining barcoded single-cell projection mapping with *in situ* detection of endogenous mRNAs exploits this unique advantage in throughput to efficiently interrogate both neuronal gene expression and long-range projections simultaneously.

To detect gene expression using BARseq2, we used a non-gap-filled padlock probe-based approach to amplify target endogenous mRNAs (Ke et al., 2013; Qian et al., 2020)(Fig. 1C, *right*). In this approach, the identity of the target is read out by sequencing a gene-identification index (GII) using Illumina sequencing chemistry *in situ*. Because the GII is a nucleotide barcode sequence that uniquely encodes the identity of a given gene, the multiplexing capacity increases exponentially as 4^N^, where N is the number of sequencing cycles. For example, a GII of length 5 can be read out with 5 sequencing cycles and can detect 4^5^ =1,024 distinct genes (although in practice a few extra cycles are used for error correction). This combinatorial coding by sequencing readout thereby allows simultaneous detection of a large number of genes using only a few cycles of imaging (Fig. 1D). Although sequencing readout offers many advantages, BARseq2 is also compatible with hybridization-based readout when necessary. The combination of non-gap-filling *in situ* sequencing of endogenous genes and the gap-filling approach for sequencing barcodes allows many genes to be detected simultaneously with projections using BARseq2.

Our goal is to combine high-throughput projection mapping with single neuron gene profiling to identify cadherin correlates of projections. In the following, we first demonstrate that, by optimizing targeted *in situ* sequencing, BARseq2 can achieve sufficient sensitivity for detection of endogenous mRNAs. We then combine *in situ* sequencing of endogenous mRNAs with *in situ* sequencing of RNA barcodes to associate the expression of cadherins with projection patterns at cellular resolution. We recapitulate previous findings of projection patterns specific to transcriptomic neuronal subtypes and identify cadherins that distinguish major projection classes. We furthermore identify a set of cadherins shared between the mouse auditory cortex and motor cortex that correlate with homologous projections within intratelencephalic (IT) neurons in both cortical areas.

### BARseq2 robustly detects endogenous mRNAs

To adequately detect genes using BARseq2, we sought to improve the detection sensitivity. In most *in situ* hybridization methods, high sensitivity is achieved by using many probes for each target mRNA (Chen et al., 2015; Codeluppi et al., 2018; Eng et al., 2019; Raj et al., 2008; Shah et al., 2016; Wang et al., 2018). We reasoned that increasing the number of padlock probes per gene might similarly improve the sensitivity of BARseq2. Indeed, we observed that tiling the whole gene with additional probes (see Methods for probe design) resulted in as much as a 46-fold increase in sensitivity compared to using a single probe (Fig. 1E; see Supplementary Note 1). Combined with other technical optimizations (Extended Data Fig. 1A, B; see Supplementary Note 1), we increased the sensitivity of BARseq2 to 60 % of RNAscope, a sensitive and commercially available FISH method (Fig. 1F; Extended Data Fig. 1C, D; see Supplementary Note 1). Hence, our optimizations allowed BARseq2 to achieve sufficiently sensitive detection of mRNAs.

To multiplex gene detection with high imaging throughput, we optimized *in situ* sequencing to robustly read out GIIs of single rolonies over many sequencing cycles. We had previously adapted Illumina sequencing chemistry to sequence neuronal somata filled abundantly with RNA barcode rolonies, i.e. DNA nanoballs generated by rolling circle amplification (Chen et al., 2019; Chen et al., 2018). However, directly applying this method to sequence single rolonies produced from individual mRNAs proved difficult due to heating cycles and harsh stripping treatments that led to loss and/or jittering of rolonies (Extended Data Fig. 1E). To allow robust sequencing of single rolonies, we optimized cryo-sectioning (see Methods) and amino-allyl dUTP concentration (Lee et al., 2014) to crosslink rolonies more extensively, achieving less spatial jitter of single rolonies between imaging cycles (Extended Data Fig. 1E-H) and stronger signals (Extended Data Fig. 1I) retained over cycles. This robust *in situ* sequencing of combinatorial GII codes allowed BARseq2 to achieve fast imaging critical for high throughput correlation of gene expression with projections.

### BARseq2 allows multiplexed detection of endogenous mRNAs *in situ*

To assess multiplexed detection of cadherins *in situ* using BARseq2, we examined the expression of 20 classical cadherins and non-clustered protocadherins expressed in the adult cortex, along with either three (in auditory cortex) or 45 (in motor cortex) cell-type markers (Fig. 2A-C). In these experiments we used up to 12 probes per gene, which resulted in sensitivity that was sufficient albeit somewhat below the maximum achievable with more probes. All but three genes were visualized using combinatorial GII codes (4-nt in auditory cortex and 7-nt in motor cortex; see Supp. Table S1); only a small subset of all possible GIIs were used, ensuring a Hamming distance of at least two bases between all pairs of GIIs in auditory cortex and three bases in motor cortex for error correction. The three remaining genes with high expression (*Slc17a7*, *Gad1*, and *Slc30a3*) were detected by hybridization. We successfully resolved and decoded 419,724 rolonies from two slices of mouse auditory cortex (1.7 mm^2^ × 10 μm per slice) and 1,445,648 rolonies from four slices of primary motor cortex (2.8 mm^2^ × 10 μm per slice). We recovered 20 rolonies in auditory cortex and 115 rolonies in motor cortex that matched two GIIs that were not used in the experiment, corresponding to an estimated error rate of 0.1 % and 0.2 %, respectively, for rolony decoding.

**Figure 2.**
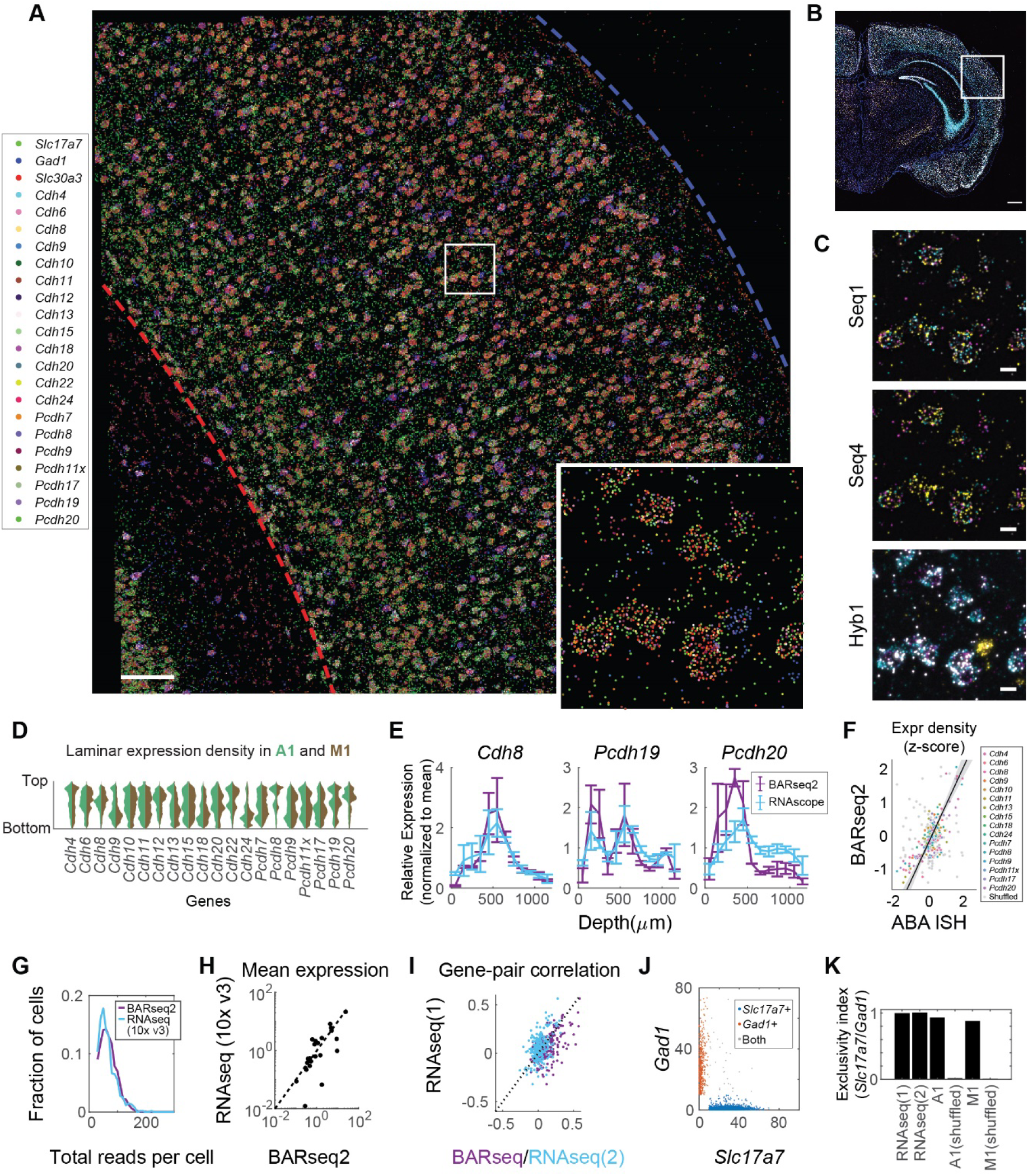
Multiplexed detection of mRNAs using BARseq2. (A) A representative image of rolonies in auditory cortex. mRNA identities are indicated on the left. The top and bottom of the cortex are indicated by the blue and red dashed lines, respectively. Scale bar = 100 μm. The inset shows a magnified view of the boxed area. (B) Low magnification image of the hybridization cycle showing the location of the area imaged in A. Scale bar = 100 μm (C) Representative images of the indicated sequencing cycle and hybridization cycle of the boxed area in A. Scale bars = 10 μm. (D) Violin plots showing the laminar distribution of cadherin expression in neuronal somata. Expression in auditory cortex and motor cortex is shown in different colors as indicated. (E) Laminar distribution of *Cdh8*, *Pcdh19*, and *Pcdh20* expression as detected by BARseq2 or FISH, normalized to the mean expression for each gene across all layers. Error bars indicate standard errors. n = 2 slices for BARseq2 and n = 3 slices for FISH. (F) Relative gene expression observed using BARseq2 (y-axis) and in Allen gene expression atlas (x-axis). Each dot represents the expression of a gene in a 100 μm bin in laminar depth. Gene identities are color-coded as indicated. Gray dots indicate correlation between data randomized across laminar positions. A linear fit and 95 % confidence intervals are shown by the diagonal line and the shaded area. (G) Distribution of total read counts per cell in BARseq2 and single-cell RNAseq using 10x v3 in auditory cortex. Only genes used in the panel detected by BARseq2 were included. (H) Mean expression for each gene detected using BARseq2 (x-axis) or single-cell RNAseq (y-axis). Each dot represents a gene. Both axes are plotted in log scale. The dotted line indicates equal expression between BARseq2 and single-cell RNAseq. (I) The correlation between pairs of genes observed in BARseq2 (x-axis, purple dots) and single-cell RNAseq (y-axis, purple dots), or in two single-cell RNAseq datasets (blue dots). The dotted line indicates that correlations between the two datasets are equal. (J) Expression of *Slc17a7* (x-axis) and *Gad1* (y-axis) in single neurons. Color codes indicate whether the neuron dominantly expressed *Slc17a7* (blue) or *Gad1* (red), or expressed both strongly (gray). (K) Exclusivity index (see Methods) of *Slc17a7* and *Gad1* in neurons in two single-cell RNAseq datasets, BARseq2 in auditory or motor cortex, and shuffled BARseq2 data.

Consistent with previous reports (Basu et al., 2015; Hayano et al., 2014; Krishna et al., 2009; Krishna et al., 2011; Lein et al., 2007; Matsunaga et al., 2015; Paul et al., 2017; Redies, 1997; Tasic et al., 2018), many cadherins were enriched in specific layers and sublayers in the cortex (Fig. 2D). Interestingly, although most cadherins had similar laminar expression in both auditory cortex and motor cortex, some cadherins were differentially expressed across the two areas. For example, *Cdh9* and *Cdh13* were enriched in L2/3 in auditory cortex, but not in motor cortex (Fig. 2D; Extended Data Fig. 2). The laminar positions of peak cadherin expression were consistent with those obtained by other methods, including RNAscope (Fig. 2E; see Supplementary Note 2) and the Allen ISH atlas (Lein et al., 2007)(Fig. 2F; Extended Data Fig. 3, Spearman correlation ρ = 0.696 comparing the gene expression density in 100 μm bins across laminae in BARseq2 and in Allen Brain Atlas; see Supplementary Note 2). Thus, BARseq2 accurately resolved the laminar expression patterns of cadherins.

We then characterized gene expression obtained by BARseq2 at single-cell resolution. We first defined excitatory and inhibitory neurons as cells having at least 10 reads of either the excitatory marker *Slc17a7* or the inhibitory marker *Gad1,* respectively. We then assigned 228,371 rolonies to 3,377 excitatory or inhibitory neurons [67.6 ± 28.8 (mean ± standard deviation) rolonies per neuron] in auditory cortex, and 752,687 rolonies to 11,492 excitatory or inhibitory neurons [65.5 ± 26.0 (mean ± standard deviation) rolonies per neuron) in motor cortex. Most cadherins showed slight differences in single-cell expression levels in these two cortical areas (Extended Data Fig. 4). In auditory cortex, the total read counts per cell was higher in BARseq2 than in single-cell RNAseq using 10x Genomics v3 (Fig. 2G; median read counts 64 for BARseq2, n = 3,337 cells compared to 57 for single-cell RNAseq, n = 640 cells, p = 5.3×10^−5^ using rank sum test). Thus, even using a limited number of probes, BARseq2 achieved sensitivity at least equal to single-cell RNAseq using 10x v3. For experiments requiring better quantification of low-expressing genes, the sensitivity could potentially be further improved by using more probes.

Further analyses showed that detection of mRNA by BARseq2 was specific. The mean expression of genes determined by BARseq2 was highly correlated with that determined by single-cell RNAseq using 10x v3 (Fig. 2H; Pearson correlation r = 0.88). Furthermore, correlations between pairs of genes in single neurons determined by BARseq2 were consistent with single-cell RNAseq using 10x v3 to a similar extent as two independent 10x v3 experiments (Fig. 2I; Pearson correlation r = 0.61 and r = 0.52 between BARseq2 and two single-cell RNAseq datasets, respectively, and r = 0.54 between the two single-cell RNAseq datasets; p = 0.78 comparing the difference in correlation between the second single-cell RNAseq dataset to either the first single-cell RNAseq or BARseq2 dataset through bootstrapping). For example, *Slc17a7* and *Gad1*, two genes expressed in either excitatory or inhibitory neurons, respectively, maintained their mutual exclusivity in both auditory cortex and motor cortex as observed by BARseq2 (Fig. 2J, K; see Supplementary Note 3). Similarly, consistent with a previous single-cell RNAseq study (Tasic et al., 2018), BARseq2 also confirmed the observation that *Slc30a3* was more highly expressed in subtypes of excitatory neurons that did not express *Cdh24* compared to projection neurons that did express *Cdh24* [Extended Data Fig. 5A, B; p = 5 × 10^−26^ using rank sum test on single-cell RNAseq data using Smart-Seq2 (n = 10,044 neurons) (Tasic et al., 2018), and p = 4 × 10^−65^ on BARseq2 data (n = 2,947 neurons)]. These results indicate that the single-cell gene expression patterns observed by BARseq2 were comparable to those of single-cell RNAseq.

Although finding cadherins that correlate with projections required only low- to medium-level of multiplexing, we wondered if BARseq2 could detect more genes in parallel, and thus be potentially useful in associating projections with larger gene panels. Because imaging time scales logarithmically with the number of genes detected (Fig. 1D), the multiplexing capacity of BARseq2 is limited not by imaging time but by potential reduction in sensitivity when more genes are probed simultaneously. To examine if multiplexing affects detection sensitivity, we probed for *Slc17a7*, *Slc30a3*, and *Gad1* either as a separate three-gene panel or as part of the 65-gene panel (20 cadherins and 45 marker genes). The mean expression density across laminar positions for the three genes were similar between the three-gene panel and the 65-gene panel (Extended Data Fig. 5C; p = 0.22 for *Slc17a7*, p = 0.49 for *Slc30a3*, and p = 0.66 for *Gad1* using rank sum tests, respectively), suggesting that targeting more genes did not affect detection sensitivity of each gene. Furthermore, the detection of the 65-gene panel in motor cortex (Fig. 3A) allowed us to classify neurons to one of nine transcriptomic neuronal types defined by single-cell RNAseq (Yao et al., 2020) (Fig. 3B; See Supplementary Note 4 and Extended Data Fig. 5D-H). Consistent with previous studies (Tasic et al., 2018; Yao et al., 2020; Zhang et al., 2020), these transcriptomic neuronal types displayed distinct laminar distributions (Fig. 3B, C; See Supplementary Note 4) and cadherin expression (Fig. 3D). These results demonstrate that BARseq2 can be applied to probe gene panels consisting of high dozens and potentially hundreds of genes, with minimal decrease in sensitivity and minimal increase in imaging time.

**Figure 3.**
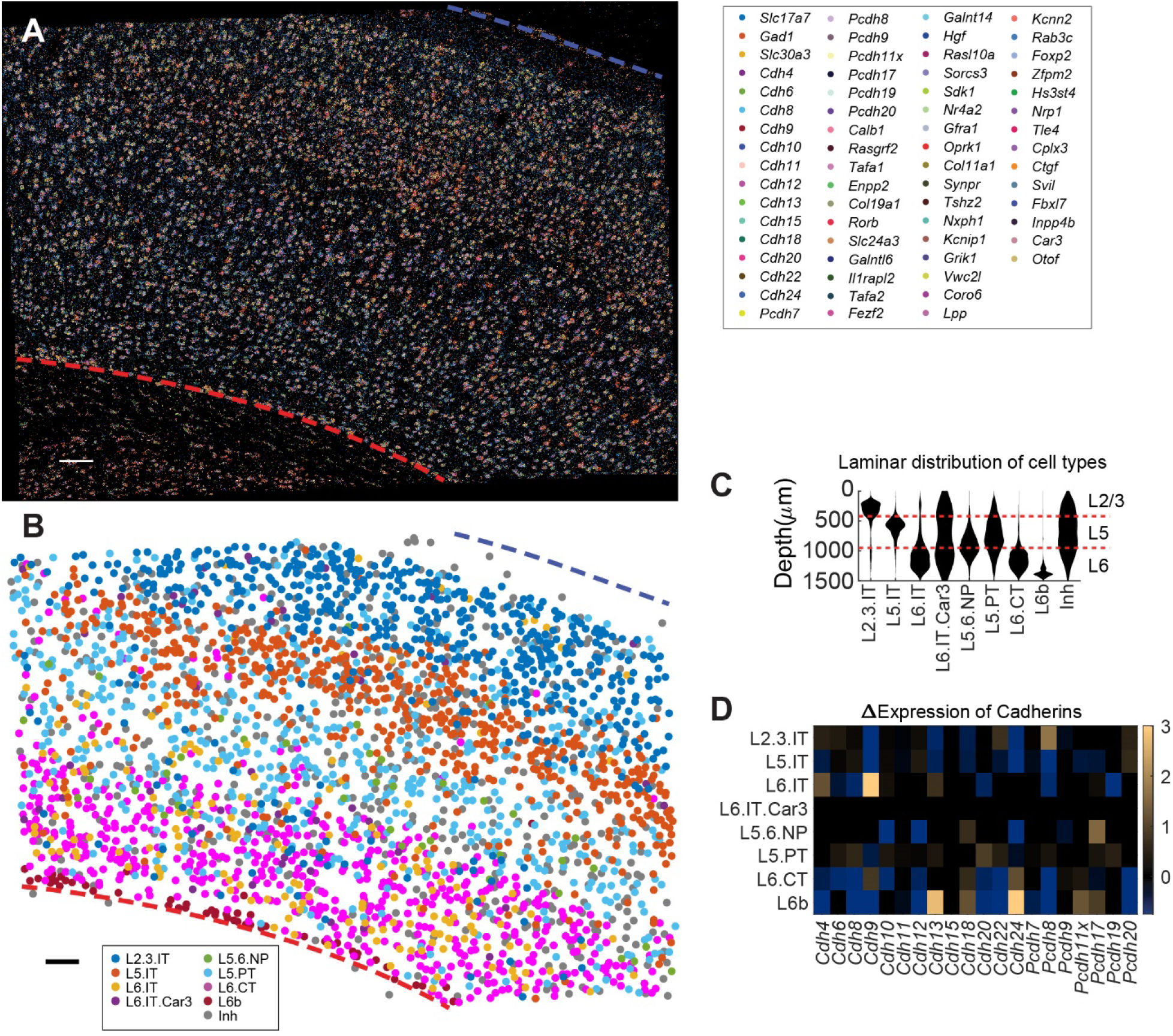
Cadherin expression across transcriptomic neuronal types in motor cortex. (A) A representative image of rolonies in motor cortex. mRNA identities are color-coded as indicated. The top and the bottom of the cortex are indicated by the blue and red dashed lines, respectively. Scale bar = 100 μm. (B) Transcriptomic cell types called based on gene expression shown in (A). (C) Laminar distribution of transcriptomic neuronal types based on marker gene expression observed by BARseq2. Layer identities are shown on the right. (D) Differential expression of cadherins across transcriptomic neuronal types identified by BARseq2. Over-expression is indicated in yellow and under-expression is indicated in blue. Only differential expression that was statistically significant was shown. Statistical significance was determined using rank sum test with Bonferroni correction for each gene between the indicated transcriptomic type and the expression of that gene across all other neuronal types.

### BARseq2 correlates gene expression to projections at cellular resolution with high throughput

To assess cadherin expression and long-range projections in the same cells, we optimized for simultaneous detection and amplification of both endogenous mRNAs and barcodes. Although both endogenous mRNAs and barcodes are amplified using padlock probe-based approaches, amplifying barcodes required the addition of a DNA polymerase to copy barcode sequences into padlock probes to allow direct sequencing of diverse barcodes (up to ~10^18^ diversity; Fig. 1C, *left*). Directly combining the two processes reduced the detection sensitivity of target mRNAs due to the addition of the DNA polymerase [Extended Data Fig. 6A; 37 ± 3 % (mean ± standard error) comparing the *Ctrl* condition to the no polymerase condition]. To preserve detection sensitivity for endogenous mRNAs while allowing the sequencing of diverse barcodes, we adjusted the concentration of the DNA polymerase to 0.001 U/μl (1/200 of the amount in the original BARseq), which doubled the sensitivity for endogenous mRNAs while also maintaining the sensitivity for barcodes (Extended Data Fig. 6A). This optimization allowed BARseq2 to detect both endogenous mRNAs and RNA barcodes together in the same neurons without compromising sensitivity.

We applied BARseq2 to study the correlation between long-range axonal projections and the expression of 20 cadherins, along with three marker genes, in motor cortex and auditory cortex in three mice. In each barcoded cell, we segmented barcoded cell bodies (Fig. 4A, *middle*) using the barcode sequencing images (Fig. 4A, *left*) and assigned rolonies amplified from endogenous genes to the segmented cells (Fig. 4A, *right*). This allowed us to map both the projection patterns of neurons (Fig. 4B, *left*) and gene expression (Fig. 4B, *right*) in the same neurons. To maintain consistency with previous studies (Chen et al., 2019)(Chen et al., unpublished observations), we sampled 11 and 35 projection targets for neurons in auditory cortex and motor cortex, respectively; these projection targets corresponded to most of the major projection targets based on bulk tracing (Oh et al., 2014). We matched barcodes in these target sites to 3,164 well-segmented barcoded neurons (1,283 from auditory cortex and 1,881 from motor cortex) from 15 slices of auditory cortex and 16 slices of motor cortex, each with 10 μm thickness. Of the barcoded neurons, 624 and 791 neurons had projections above the noise floor in auditory cortex and motor cortex, respectively (Fig. 4C). Most neurons [53 % (329/624) in auditory and 89 % (703/791) in motor cortex] projected to multiple brain areas. These observations were largely consistent with previous BARseq experiments in auditory and motor cortex performed without assessing gene expression (Chen et al., 2019)(Chen et al., unpublished observations), confirming that the modifications for BARseq2 did not compromise projection mapping.

**Figure 4.**
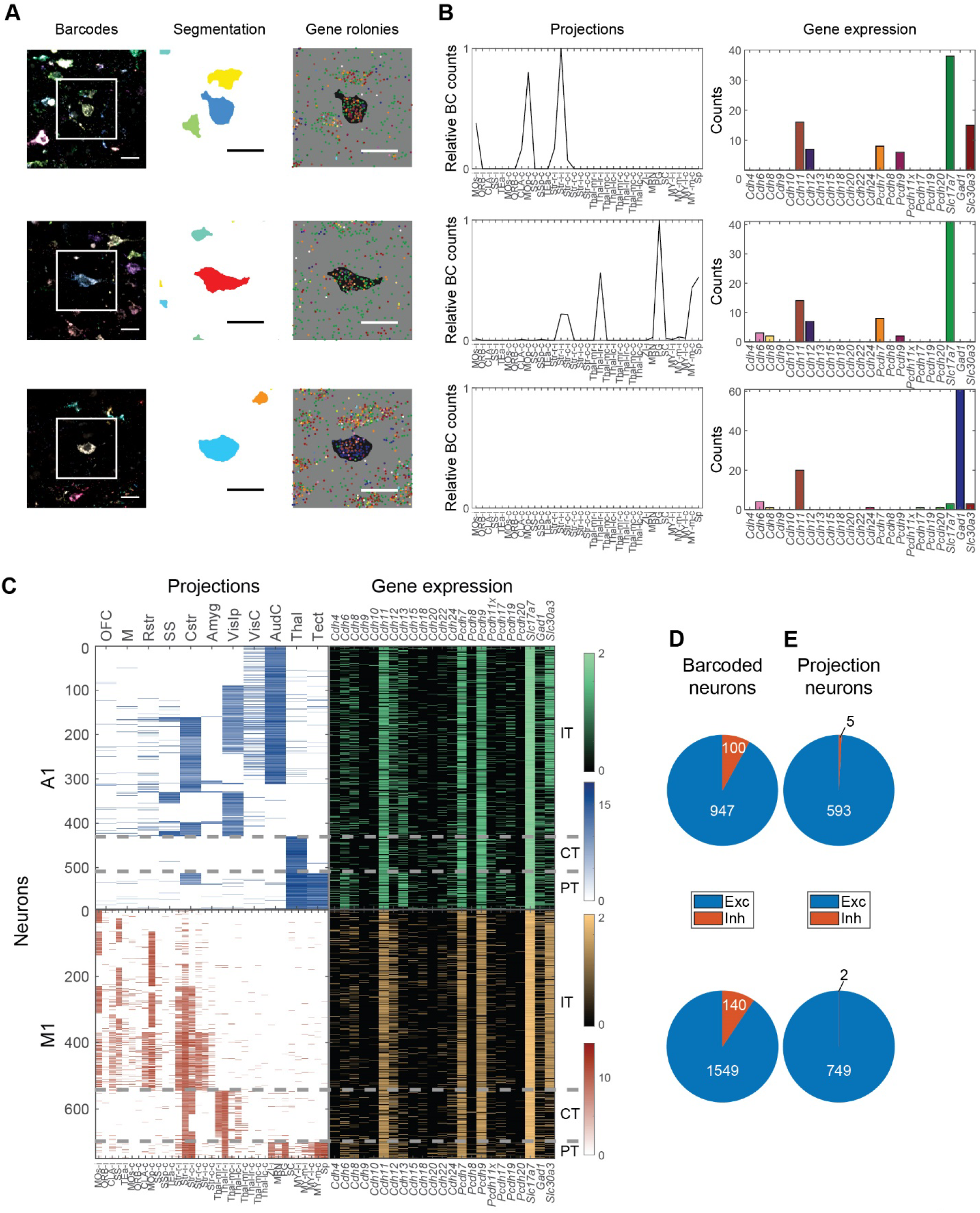
Correlating gene expression to projections using BARseq2. (A) False-colored barcode sequencing images (*left*), soma segmentations (*middle*), and gene rolonies (*right*) of three representative neurons from the motor cortex. The segmentation and gene rolony images correspond to the white squared area in the barcode images. In the gene rolony images, the areas corresponding to the soma segmentations of the target neurons are in black. All scale bars = 20 μm. (B) Projections (*left*) and gene expression (*right*) of the target neurons shown in (A). The bars indicating gene expression are colored using the same color code as that in the gene rolony plots in (A). The neurons shown in the first two rows are excitatory projection neurons, whereas the neuron shown in the bottom row is an inhibitory neuron without projections. See Supp. Table S2 for the brain areas corresponding to each abbreviated target area. Projections (*left*) and gene expression (*right*) of neurons in auditory cortex (*top*) and motor cortex (*bottom*). Each row represents a barcoded projection neuron. Both projections and gene expression are shown in log scale. Major projection neuron classes determined by projection patterns are indicated on the right. (D) (E) The number of excitatory neurons (blue) or inhibitory neurons (red) in all barcoded neurons or barcoded projection neurons (E). Neurons in auditory cortex are shown in the top row and those in motor cortex are shown in the bottom row.

### BARseq2 recapitulates known projection biases across transcriptomic cell types

Although we have demonstrated that BARseq2 can read out cadherin expression and projections in the same neurons, one might be concerned that barcoding neurons using Sindbis virus could disrupt gene expression (Fros and Pijlman, 2016). To find the relationship between genes and projections, one would require that the gene-gene relationship in Sindbis-infected single neurons reflects that in non-infected neurons, but changes in absolute gene expression level would have little effect. Reassuringly, previous reports have shown that the relationship among genes in single neurons is indeed largely preserved despite a reduction in the absolute expression of genes in Sindbis-infected cells (Chen et al., 2019; Klingler et al., 2018). Furthermore, correlations between transcriptomic types and projections revealed in Sindbis-infected neurons were corroborated by other methods that did not require Sindbis infection (Chen et al., 2019; Wang et al., 2019). Consistent with these previous reports, we observed that the correlations between pairs of genes in the barcoded neurons were consistent with those in non-barcoded neurons (Extended Data Fig. 6B). For example, the expression of the excitatory marker *Slc17a7* and the inhibitory marker *Gad1* remained mutually exclusive in barcoded neurons in both auditory cortex and motor cortex (Extended Data Fig. 6C, D). This mutual exclusivity was preserved despite an overall reduction in mRNA expression (Extended Data Fig. 6E; median reads of 38 in barcoded cells in both auditory and motor cortex, compared to 64 and 48 in non-barcoded cells in the two cortical areas, respectively). Similarly, *Slc30a3* remained differentially expressed across barcoded excitatory neurons with or without *Cdh24* expression as it was in non-barcoded excitatory neurons (Extended Data Fig. 6F; p = 1 × 10^−6^ using rank sum test, n = 810 neurons). Although our observations cannot rule out the possibility that a small subset of genes may be disrupted by Sindbis infection (e.g. viral response genes), these results suggest that the co-expression relationships of most genes in Sindbis-infected neurons reflect those in non-infected cells. Therefore, the relationship between gene expression and projections resolved by BARseq2 likely reflects that in non-barcoded neurons.

To further test whether BARseq2 can capture the relationship between gene expression and projections, we asked if we could identify differences in projection patterns across transcriptomic neuronal types that could also be validated by previous studies and/or other experimental techniques. We performed these analyses at three different levels of granularity: between excitatory neurons and inhibitory neurons, among major classes of excitatory neurons, and among transcriptomic subtypes of intratelencephalic (IT) neurons in auditory cortex.

First, BARseq2 confirmed that most barcoded neurons with long-range projections were excitatory, not inhibitory. To distinguish between excitatory and inhibitory neurons, we categorized a neuron as excitatory or inhibitory if (1) the neuron had higher expression of the excitatory marker *Slc17a7* or the inhibitory marker *Gad1*, respectively; and (2) the marker was expressed at greater than five reads in the cell. This threshold resulted in 2,496 excitatory neurons (947 in auditory and 1,549 in motor cortex) and 240 inhibitory neurons (100 in auditory cortex and 140 in motor cortex) (Fig. 4D). Consistent with the fact that the majority of long-range projection neurons in the cortex are excitatory, 1,342 of 2,501 (54 %) excitatory neurons, including 593 in auditory cortex and 749 in motor cortex, had detectable projections, whereas only 7 of 240 (3 %) inhibitory neurons (5 in auditory cortex and 2 in motor cortex) had detectable projections (Fig. 4E; see Supplementary Note 5 and Extended Data Fig. 6G, H). The excitatory neurons that did not have detectable projections were likely neurons that projected only locally or to unsampled nearby cortical areas (see Supplementary Note 5). Hence, BARseq2 accurately observed the fact that projection neurons in the cortex are predominantly excitatory and express the excitatory marker *Slc17a7*, not the inhibitory marker *Gad1*.

Second, BARseq2 revealed differential gene expression across major classes of neurons defined by projections. We found that many cadherins (8 for auditory cortex and 12 for motor cortex) were differentially expressed across intratelencephalic (IT) neurons, pyramidal tract (PT) neurons, and corticothalamic (CT) neurons that were defined by projections (Harris and Shepherd, 2015) (Fig. 5A-C; see Methods for the classification of neurons to projection classes). Several cadherins were consistently differentially expressed in both cortical areas. For example, *Cdh6* and *Cdh13* were over-expressed in PT neurons compared to the other two classes, whereas *Cdh8* was under-expressed in CT neurons compared to the other two classes (FDR < 0.05 using rank sum test). In addition, we also found nine cadherins that were differentially expressed across the two cortical areas in at least one class (Fig. 5D; FDR < 0.05 using rank sum tests). Since IT, PT, and CT neurons can also be classified based on transcriptomic data alone, we compared our findings to the expression of these cadherins observed by single-cell RNAseq (Tasic et al., 2018). The differences in cadherin expression across pairs of classes identified by BARseq were consistent with those observed by single-cell RNAseq (Extended Data Fig. 7A; the rank correlation of the differences in cadherin expression across major neuronal types was 0.61 between BARseq and single-cell RNAseq, compared to 0.39 between auditory and motor cortex in BARseq; see Supplementary Note 6). Thus, BARseq2 identified cadherin correlates of major neuronal classes that were consistent with those observed using single-cell RNAseq.

**Figure 5.**
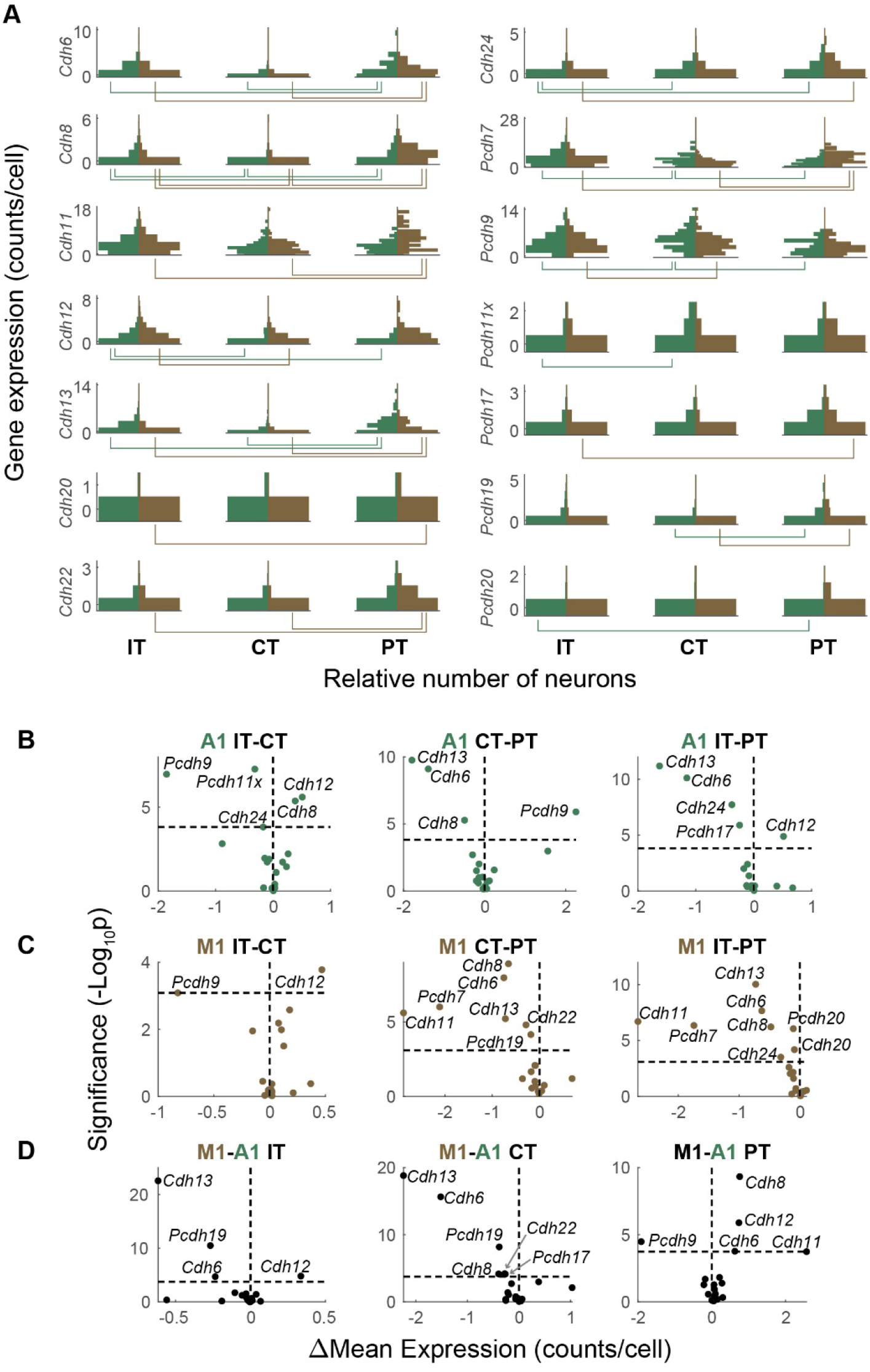
Differential cadherin expression across major classes and cortical areas. (A) Vertical histograms of the expression (raw counts per cell) of cadherins that were differentially expressed across major classes in either auditory or motor cortex. Y-axes indicate gene expression level (counts per cell) and x-axes indicate number of neurons at that expression level. The numbers of neurons are normalized across plots so that the bins with the maximum number of neurons have equal bar lengths. Gene expression in auditory cortex (green) are shown on the left in each plot, and gene expression in motor cortex (brown) are shown on the right in each plot. Lines beneath each plot indicate pairs of major classes with different expression of the gene (FDR < 0.05). (B)(C) Volcano plots of cadherins that were differentially expressed across pairs of major classes in auditory cortex (B) or motor cortex (C). Y-axes indicate significance and x-axes indicate effect size. The horizontal dashed lines indicate significance level for FDR < 0.05, and the vertical dashed lines indicate equal expression. (D) Volcano plots of cadherins that were differentially expressed across auditory and motor cortex in the indicated major classes. Y-axes indicate significance and x-axes indicate effect size. The horizontal dashed lines indicate significance level for FDR < 0.05, and the vertical dashed lines indicate equal expression.

Finally, BARseq2 confirmed known biases in projection patterns across transcriptomic IT subtypes in auditory cortex (Extended Data Fig. 7B, C). Previous studies using both barcoding-based strategy and single-cell tracing have identified distinctive projection patterns for two transcriptomic subtypes of IT neurons, IT3 (L6 IT) and IT4 (L6 *Car3*+ IT) (Chen et al., 2019; Wang et al., 2019). To test if we could capture the same projection specificity of transcriptomic subtypes, we mapped projection patterns to projection clusters identified in a previous study in auditory cortex, and used a combination of gene expression and laminar position to distinguish four transcriptomic subtypes of IT neurons (Chen et al., 2019)(see Methods for detailed definitions). As expected, the two transcriptomic subtypes (IT3 and IT4) predominantly found in L5 and L6 were indeed more likely to project only to the ipsilateral cortex, without projections to the contralateral cortex or the striatum (p = 4 ×10^−7^ comparing the fraction of neurons with only ipsilateral cortical projections in IT3/IT4 to the fraction of them in IT1/IT2 using Fisher’s test; Extended Data Fig. 7B, C). Between IT3 and IT4, IT4 neurons were more likely to project ipsilaterally (58 % IT3 neurons compared to 92 % IT4 neurons, p = 1×10^−4^ using Fisher’s test), whereas IT3 neurons were more likely to project contralaterally (66 % IT3 neurons compared to 14 % IT4 neurons, p = 5 ×10^−8^ using Fisher’s test). Thus, BARseq2 recapitulated known projection differences across transcriptomic subtypes of IT neurons.

### BARseq2 identifies cadherin correlates of IT projections

Having established that BARseq2 identified gene correlates of projections that were consistent with previous studies, we then moved to identify cadherins that correlate with projections within IT neurons. We first grouped projections to different areas based on the likelihood of co-innervation, and then identified genes that correlated with these groups of projections. The projections of IT neurons to multiple brain areas correlated with each other in both auditory cortex and motor cortex (Chen et al., 2019)(Fig. 6A). For example, neurons in the auditory cortex that projected to the somatosensory cortex (SS) were also more likely to project to the ipsilateral visual cortex (VisIp), but not the contralateral auditory cortex (AudC). To exploit these correlations, we used non-negative matrix factorization (NMF)(Lee and Seung, 1999), an algorithm related to PCA but with the added constraint that projections be non-negative, to represent the projection pattern of each neuron as the sum of several “projection modules.” Each of these modules (six modules for the motor cortex and three for the auditory cortex; Fig. 6B) consisted of subsets of projections that were likely to co-occur. We named these modules by the main projections (cortex, CTX, or striatum, STR) followed by the side of the projection (ipsilateral, −I, or contralateral, −C). For some modules, we further indicated that the projections were to the caudal part of the structure by prefixing with “C” (e.g. CSTR-I or CCTX-I). A small number of projection modules could explain most of the variance in projections (three modules and six modules explained 84 % and 87 % of the variance in projections to nine areas in auditory cortex and 18 areas in motor cortex that IT neurons project to, respectively; Fig. 6C).

**Figure 6.**
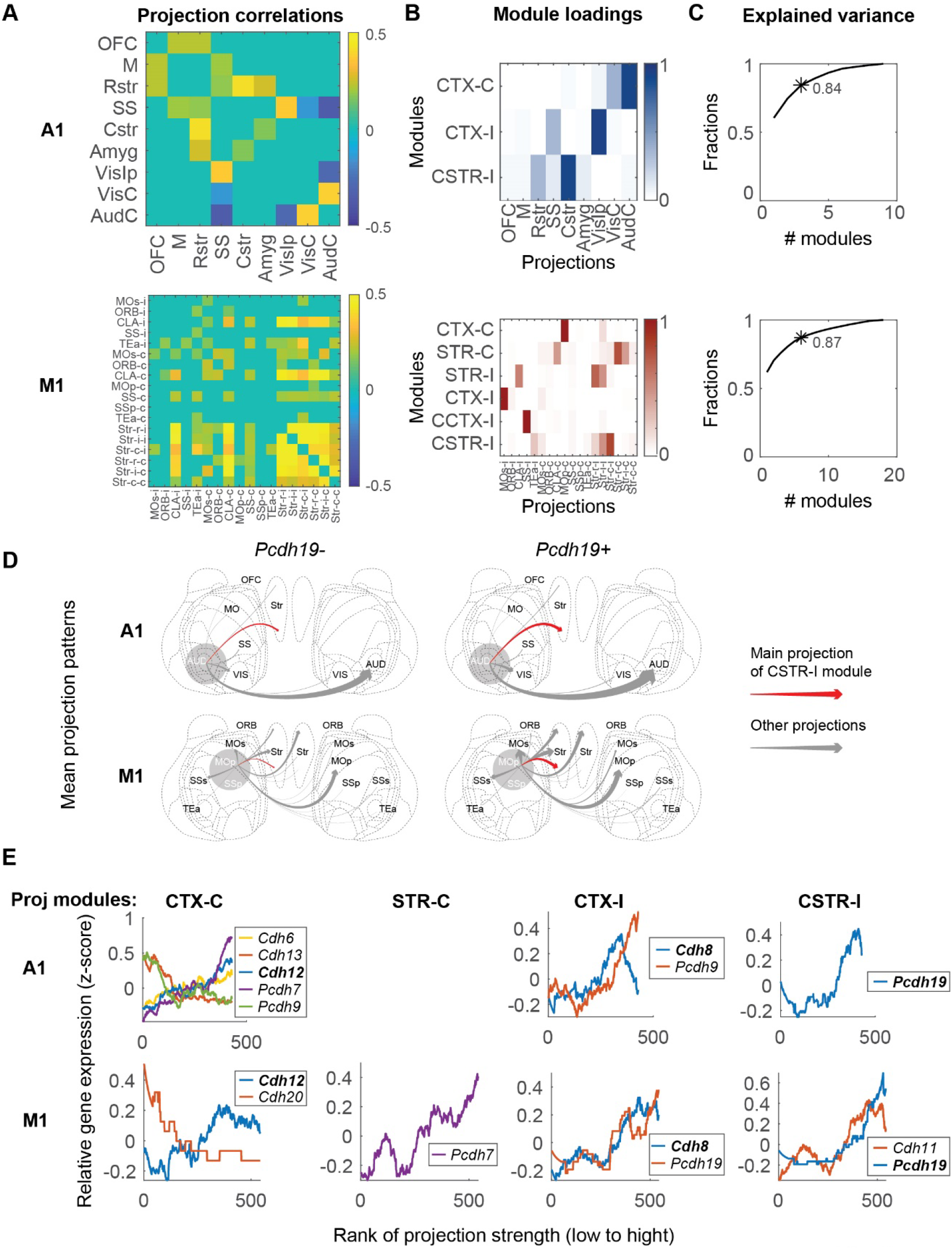
Cadherins correlate with diverse projections of IT neurons. (A) Correlation of projections to different brain areas in IT neurons of auditory cortex (*top*) or motor cortex (*bottom*). Only significant correlations are shown. (B) Projection modules of IT neurons in auditory cortex (*top*) or motor cortex (*bottom*). Each row represents a projection module. Columns indicate projections to different brain areas. (C) The fractions of variance explained by different numbers of projection modules in auditory cortex (*top*) and motor cortex (*bottom*). The numbers of projection modules that correspond to those in (B) are labeled with an asterisk with the fraction of variance explained indicated. (D) Mean projection patterns of neurons in A1 (*top*) and M1 (*bottom*) with or without *Pcdh19* expression. The thickness of arrows indicates projection strength (barcode counts). Red arrows indicate projections that correspond to the strongest projection in the CSTR-I projection modules. (E) The expression of cadherins (y-axes) that were rank correlated with the indicated projection modules in auditory cortex (*top row*) and motor cortex (*bottom row*). Neurons (x-axes) are sorted by the strengths of the indicated projection modules. Only genes that were significantly correlated with projection modules are shown (FDR < 0.1). Genes that were correlated with the same projection modules in both areas are shown in bold.

We found many cadherins whose expression co-varied with projection modules (Supp. Fig. S1). For example, auditory cortex neurons expressing *Pcdh19* were stronger in the CSTR-I projection module than those not expressing *Pcdh19* [Fig. 6D, *top*; p = 5 × 10^−4^ comparing the CSTR-I module in neurons with (n = 83) or without (n = 346) *Pcdh19* expression using rank sum test]. Surprisingly, the same association between *Pcdh19* and the CSTR-I projection module was also seen in motor cortex (Fig. 6D, *bottom*; p = 4 × 10^−6^ using rank sum test, n = 31 for *Pcdh19+* neurons and n = 512 for *Pcdh19-* neurons), despite the overall differences in projections from these two areas. Similarly, *Cdh8* was correlated with the CTX-I module and *Cdh12* was correlated with the CTX-C module (Fig. 6E, FDR < 0.1) in both auditory and motor cortex. These correlations were independently validated by retrograde tracing using cholera toxin subunit B (CTB) and FISH (Extended Data Fig. 8A-E; See Supplementary Note 7). Additional cadherins, including *Cdh6*, *Cdh11*, *Cdh20*, *Pcdh7*, and *Pcdh9*, were also correlated with projection modules in at least one of the two areas (Fig. 6E, FDR < 0.1; Supp. Fig. S1). Our observations that the same cadherins correlated with similar projection modules in both areas suggest that a common molecular logic might underscore the organization of projections across cortical areas beyond class-level divisions.

Although gene expression is inherently noisy, the expression of many genes is correlated in single neurons. We reasoned that the correlations among genes might allow us to identify additional relationships between gene expression and projections that were missed by analyzing each gene separately. To exploit the correlations among genes, we grouped 16 cadherins into three meta-analytic co-expression modules based on seven single-cell RNAseq datasets of IT neurons in motor cortex (Fig. 7A; Extended Data Fig. 9A, B) (Yao et al., 2020). To obtain the modules, we followed the rank-based network aggregation procedure defined by Ballouz et al. (2015) and Crow et al. (2016) to combine the seven dataset-specific gene-gene co-expression networks into an aggregated network, and then grouped together genes showing consistent excess correlation using the dynamic cutting tree algorithm (Langfelder et al., 2008). Two co-expressed modules were associated with projections: Module 1 was associated with contralateral striatal projections (STR-C projection module), and Module 2 was associated with ipsilateral caudal striatal projections (CSTR-I; Fig. 7B, C; Extended Data Fig. 9C, D). These associations between the co-expression modules and projections were consistent with, but stronger than, associations between individual genes contained in each module and the same projections (Extended Data Fig. 9E). Interestingly, these co-expression modules were enriched in multiple transcriptomic subtypes of IT neurons, but these transcriptomic subtypes were found in multiple branches of the transcriptomic taxonomy (Fig. 7D; Extended Data Fig. 9F). For example, Module 1 is associated with transcriptomic subtypes of IT neurons in L2/3, L5, and L6. This result is consistent with previous observations (Chen et al., 2019; Tasic et al., 2018; Zhang et al., 2020) that first-tier transcriptomic subtypes of IT neurons (i.e. subtypes of the highest level in the transcriptomic taxonomy within the IT class) appeared to share projection patterns, and further raises the possibility that transcriptomic taxonomy does not necessarily capture differences in projections. Taken together, our finding that projections correlate with cadherin co-expression modules independent of transcriptomic subtypes demonstrates that BARseq2 can reveal intricate relationships between gene expression and projection patterns.

**Figure 7.**
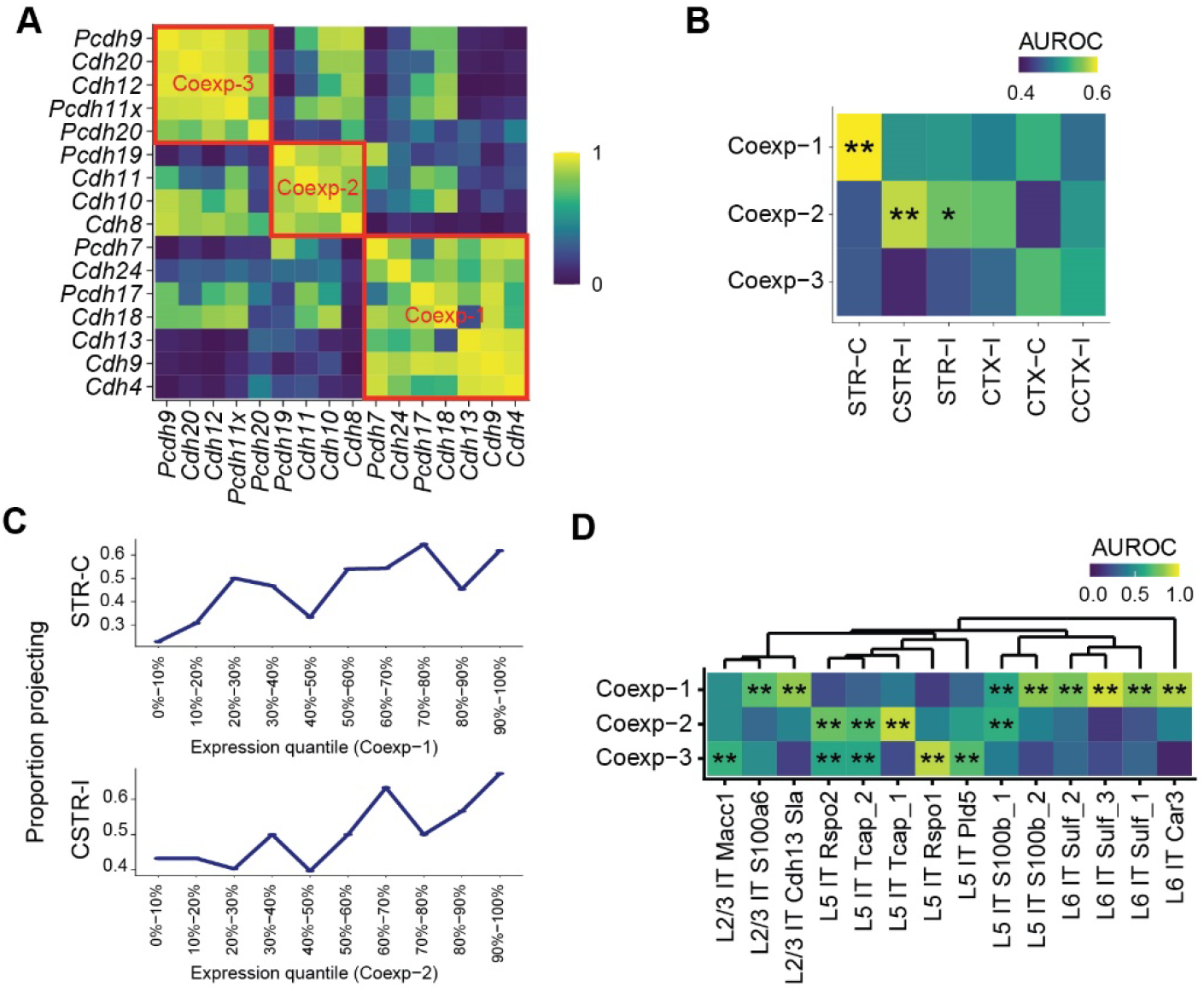
Gene co-expression modules correlate with diverse projections of IT neurons. (A) Correlation among cadherins as identified using single-cell RNAseq in IT neurons in motor cortex (Yao et al., 2020). Three co-expression modules are marked by red squares. Cadherins that did not belong to any module were not shown. (B) Association between cadherin co-expression modules and projection modules (AUROC). Significant associations are marked by asterisks (*FDR < 0.1, **FDR < 0.05). (C) Fractions of neurons with the indicated projection modules as a function of co-expression module expression. Neurons are binned by gene module quantiles as indicated. (D) Association of the three co-expression modules in transcriptomic IT neurons in the scSS dataset (AUROC, significance shown as in B).

## Discussion

BARseq2 combines high-throughput mapping of projections to many brain areas with multiplexed detection of gene expression at single-cell resolution. Because BARseq2 is high-throughput, we are able to correlate gene expression and projection patterns of thousands of individual neurons in a single experiment, and thereby achieve statistical power that would be challenging to obtain using other single-cell techniques. By applying BARseq2 to two distant cortical areas—primary motor and auditory cortex—in the adult mouse, we identified cadherin correlates of diverse projections. Our results suggest that BARseq2 provides a path to discovering general organization of gene expression and projections that are shared across the cortex.

### BARseq2 detects multiplexed gene expression with high throughput

To correlate panels of genes, such as cadherins, to projections, we designed BARseq2 to detect gene expression with high throughput, multiplex to dozens of genes, have sufficient sensitivity, and be compatible with barcoding-based projection mapping. To satisfy these needs, we based BARseq2 on padlock probe-based approaches (Ke et al., 2013; Qian et al., 2020). With additional optimizations for sensitivity, sequencing readout, and compatibility with barcode sequencing, we successfully used BARseq2 to identify cadherins that correlate with projections.

One of the critical requirements for BARseq2 is high throughput when reading out many genes. Through strong amplification of mRNAs, combinatorial coding, and robust readout using Illumina sequencing chemistry (Chen et al., 2019; Chen et al., 2018), BARseq2 achieves fast imaging at low optical resolution compared to many other imaging-based spatial transcriptomic methods (Chen et al., 2015; Codeluppi et al., 2018; Eng et al., 2019; Shah et al., 2016; Zhang et al., 2020). Further optimizations, including computational approaches for resolving spatially mixed rolonies (Chen et al., 2020), have the potential to increase imaging throughput even further.

Another critical optimization was increasing the low sensitivity that early versions of the padlock probe-based technique suffered from, unless special and expensive primers were used (Ke et al., 2013). Inspired by other spatial transcriptomic methods, we and others (Qian et al., 2020) have found that tiling target genes with multiple probes could greatly improve the sensitivity. This design allowed variable sensitivity for different experimental purposes. Although we identified cadherin correlates of projections using a modest number of probes per gene to achieve sensitivity similar to single-cell RNAseq using 10x Genomics v3, we could achieve much higher sensitivity using more probes. This high and tunable sensitivity, combined with the fact that the gene multiplexing capacity of BARseq2 is not limited by imaging time, opens potential applications of BARseq2 to a wide range of questions that require high-throughput interrogation of gene expression *in situ*.

### BARseq2 reveals gene correlates of projections

BARseq2 exploits the high-throughput axonal projection mapping that BARseq offers to identify gene correlates of diverse projections. BARseq has sensitivity comparable to single neuron tracing (Han et al., 2018). Although the spatial resolution of BARseq for projections is lower than conventional single neuron tracing, it offers throughput that is several orders of magnitude higher than the state-of-the-art single-cell tracing techniques (Lin et al., 2018; Winnubst et al., 2019). This high throughput allows BARseq to reveal higher-order statistical structure in projection patterns that would have been difficult to observe using existing techniques, such as single-cell tracing (Chen et al., 2019; Kebschull et al., 2016). The increased statistical power of BARseq, obtained at the cost of some spatial resolution, is reminiscent of different clustering power across single-cell RNAseq techniques of varying throughput and read depth (Ding et al., 2020; Yao et al., 2020). The high throughput of BARseq thereby provides a powerful asset for investigating the organization of projection patterns and their relationship to gene expression.

BARseq2 enables simultaneous measurement of multiplexed gene expression and axonal projections to many brain areas, at single neuron resolution and at a scale that would be difficult to achieve with other approaches. For example, *Cre*-dependent labeling allows interrogation of the gene expression and projection patterns of a genetically defined subpopulation of neurons (Chen et al., 2019). However, this approach lacks cellular resolution and is limited by the availability of *Cre* lines and requires that a neuronal population of interest be specifically distinguished by the expression of one or two genes. The combination of single-cell transcriptomic techniques with retrograde labeling does provide cellular resolution, but can only interrogate projections to one or at most a small number of brain areas at a time (Economo et al., 2018; Kim et al., 2019; Tasic et al., 2018; Zhang et al., 2020). The inability to interrogate projections to many brain areas from the same neuron would miss higher-order statistical structures in projections, which are non-random (Han et al., 2018) and provide additional information regarding other properties of the neurons, such as laminar position and gene expression (Chen et al., 2019)(Chen et al., unpublished observations). The projections of individual neurons to multiple brain areas can be obtained using multiplexed single-cell tracing (Lin et al., 2018; Wang et al., 2019; Winnubst et al., 2019), but the throughput of these methods remains relatively low. Moreover, many advanced single-cell tracing techniques require special sample processing that hinders multiplexed interrogation of gene expression in the same sample. BARseq2 thus provides a powerful tool for probing the relationships between gene expression and projection patterns.

### Cadherins correlate with diverse projections of IT neurons

Using BARseq2, we identified several cadherins that correlate with homologous IT projections in both auditory and motor cortex, two spatially and transcriptomically distant areas with distinct cortical and subcortical projection targets. We speculate that these cadherin correlates of projections shared across areas may represent the remnants of a common developmental program that establishes similar projections (Custo Greig et al., 2013), or may be needed for ongoing functions or maintenance of projections. Our findings raise the possibility that a shared molecular code might underlie the diversity of cortical projections.

Although the cadherin correlates of projections were observed in adult neurons and thus did not necessarily reflect the genes that specified projections during development, our observations are compatible with the “cadherin code hypothesis” (Duan et al., 2014; Duan et al., 2018; Krishna et al., 2011; Redies, 1997), according to which combinations of cadherins can specify complex projection patterns. Although recent studies have provided evidence of cadherins in specifying diverse connectivity in the retina (Duan et al., 2014; Duan et al., 2018), this hypothesis remains difficult to test systematically in more complex and heterogeneous circuits such as the cortex due to a lack of high throughput techniques with high multiplexing capacity for gene detection. BARseq2 possesses the high throughput and multiplexing qualities essential for investigating such questions, and could potentially be applied to circuit development to illuminate genetic and anatomical changes over developmental trajectories. BARseq2 thus provides a path to discovering the myriad genetic programs that specify and/or correlate with long-range projections in both developing and mature animals.

### BARseq2 builds a unified description of neuronal diversity

Recent developments in methodologies have enabled the high-throughput interrogation of individual neuronal properties, such as connectivity, neuronal activity, genomic signatures, and developmental lineage, at cellular resolution. However, a comprehensive understanding of neuronal circuits further requires the examination of how distinct neuronal characteristics relate to each other. It is essential to correlate different neuronal properties at cellular resolution to further our understanding of neuronal circuitry and function.

BARseq2 represents an important step toward this goal by integrating multiplexed gene expression and neuronal projections to many brain areas at cellular resolution with high throughput. Although we focused on the relationship between cadherin expression and projections, our results suggest a complex relationship between transcriptomic cell types and projections: the cadherin co-expression modules that correlated with projections were enriched in multiple transcriptomic clusters across different branches of the transcriptomic taxonomy. This observation raises the interesting possibility that the hierarchy of transcriptomic cell types does not necessarily capture distinctions in long-range connectivity, a hypothesis that is consistent with several previous studies (Chen et al., 2019; Tasic et al., 2018; Zhang et al., 2020).

Because BARseq2 integrates neuronal properties using spatial information, it is potentially compatible with other *in situ* assays, such as immunohistochemistry, two-photon calcium imaging, and dendritic morphological reconstruction. BARseq2 can additionally characterize genetic labeling tools, such as *Cre*-lines (Chen et al., 2019), to potentially allow genetic access and manipulation of neuronal subpopulations. Interrogating and manipulating neuronal populations in the context of various neuronal properties with high throughput and cellular resolution can elucidate the fundamental rules that govern organization of cortical neuronal diversity. By spatially correlating various neuronal properties in single neurons, BARseq2 represents a feasible path towards achieving a comprehensive description of neuronal circuits.

## Extended data figure legends

**Extended Data Figure 1.**
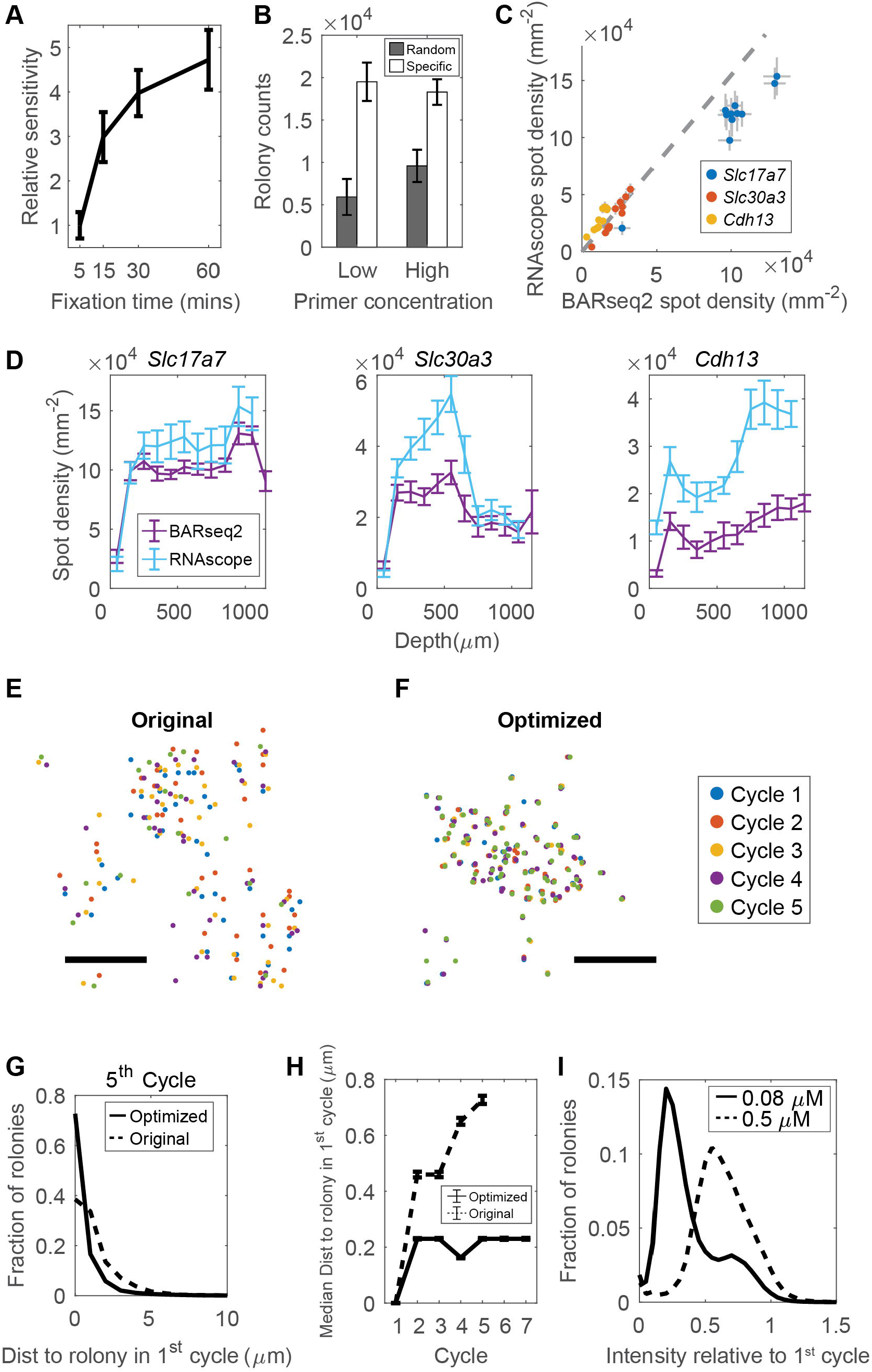
Optimization of BARseq2 for detecting endogenous mRNAs. (A) Relative sensitivity (mean ± standard error, y-axis) of BARseq2 in detecting *Slc17a7* using the indicated fixation times (x-axis). The sensitivity is normalized to that achieved with 5 mins of fixation. n = 4 slices for each time point. (B) Rolony counts for *Slc17a7* using either random primers (gray) or specific primers (white) at two different concentrations. For random primers, the two concentrations used were 5 μM (low) and 50 μM (high). For specific primers, the two concentrations were 0.5 μM (low) and 5 μM (high). Error bars indicate standard errors. n = 2 slices for each condition. (C) (D) Quantification of BARseq2 sensitivity compared to RNAscope. (C) Spot density detected by BARseq2 (x-axis) or RNAscope (y-axis) in each 100 μm bin along the laminar axis in auditory cortex. Error bars indicate standard errors. The dashed line indicates linear fit for *Slc30a3* and *Cdh13*. Slope = 1.65 and R^2^ = 0.73. n = 5 slices for both BARseq2 and RNAscope. (D) Laminar distribution of the indicated genes detected by BARseq2 and RNAscope. Error bars indicate standard errors. The data used were the same as in (C). (E)(F) Positions of rolonies across five cycles of sequencing using the original BARseq protocol (E) or the optimized BARseq2 protocol (F). Scale bars = 10 μm. The sequencing cycles in which the rolonies were imaged are color-coded as shown on the right. (G) The distribution of minimum distance between rolonies imaged in the first cycle and in the fifth cycle using the original or the optimized protocol. (H) Median distance between rolonies imaged in the indicated cycles (x-axis) and the closest rolonies imaged in the first cycle using the original or the optimized protocol. Error bars indicate standard errors. (I) The distribution of rolony intensities after 6 sequencing cycles and one stripping step, normalized to the intensities in the first sequencing cycle. Amino-allyl dUTP concentrations used are indicated. n = 128,976 rolonies for 0.08 μM and n = 113,235 rolonies for 0.5 μM.

**Extended Data Figure 2.**
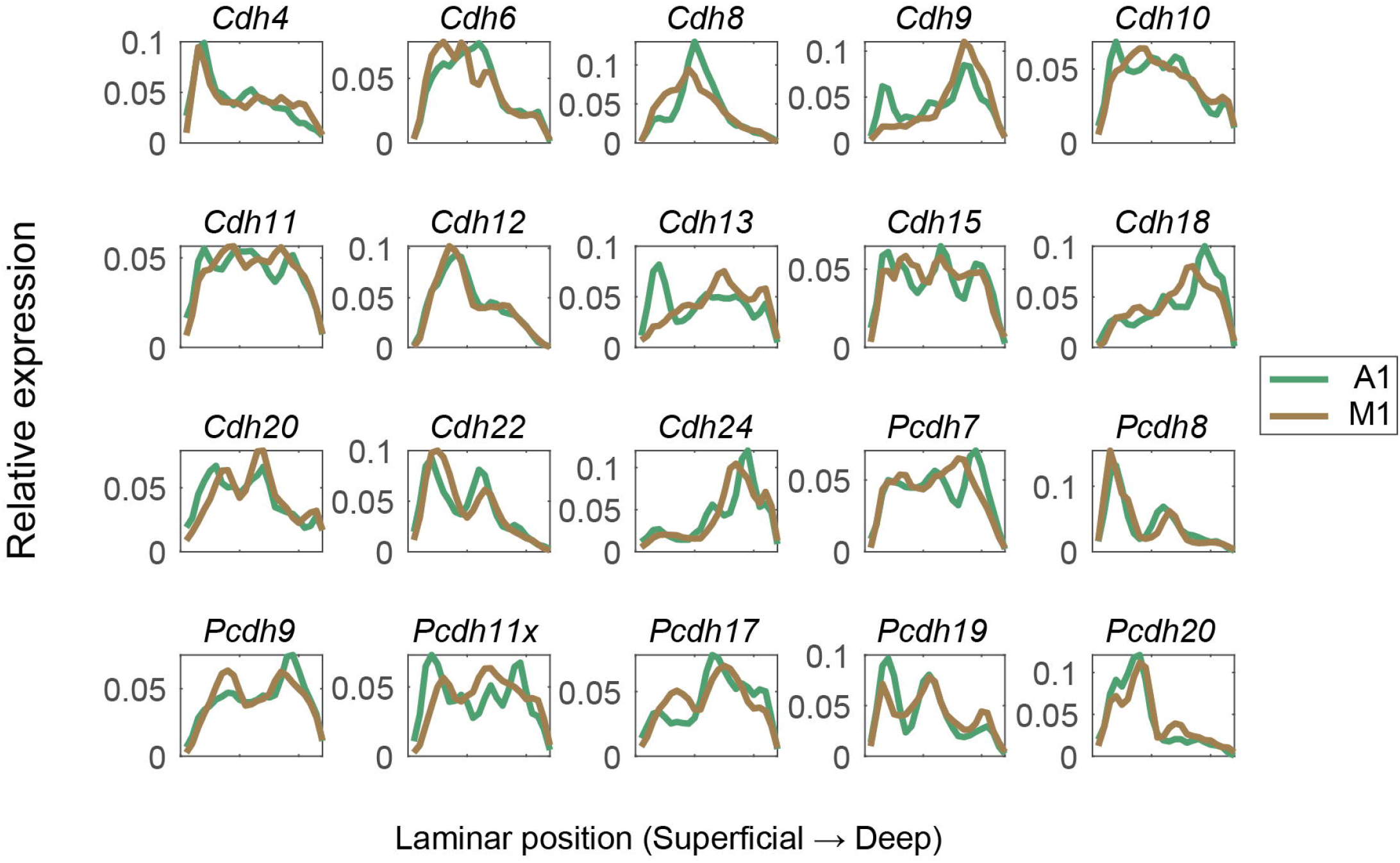
Laminar distribution of cadherins in auditory cortex (green) and motor cortex (brown). In both cortical areas, cortical depth is normalized so that the bottom and the top of the cortex match between M1 and A1.

**Extended Data Figure 3.**
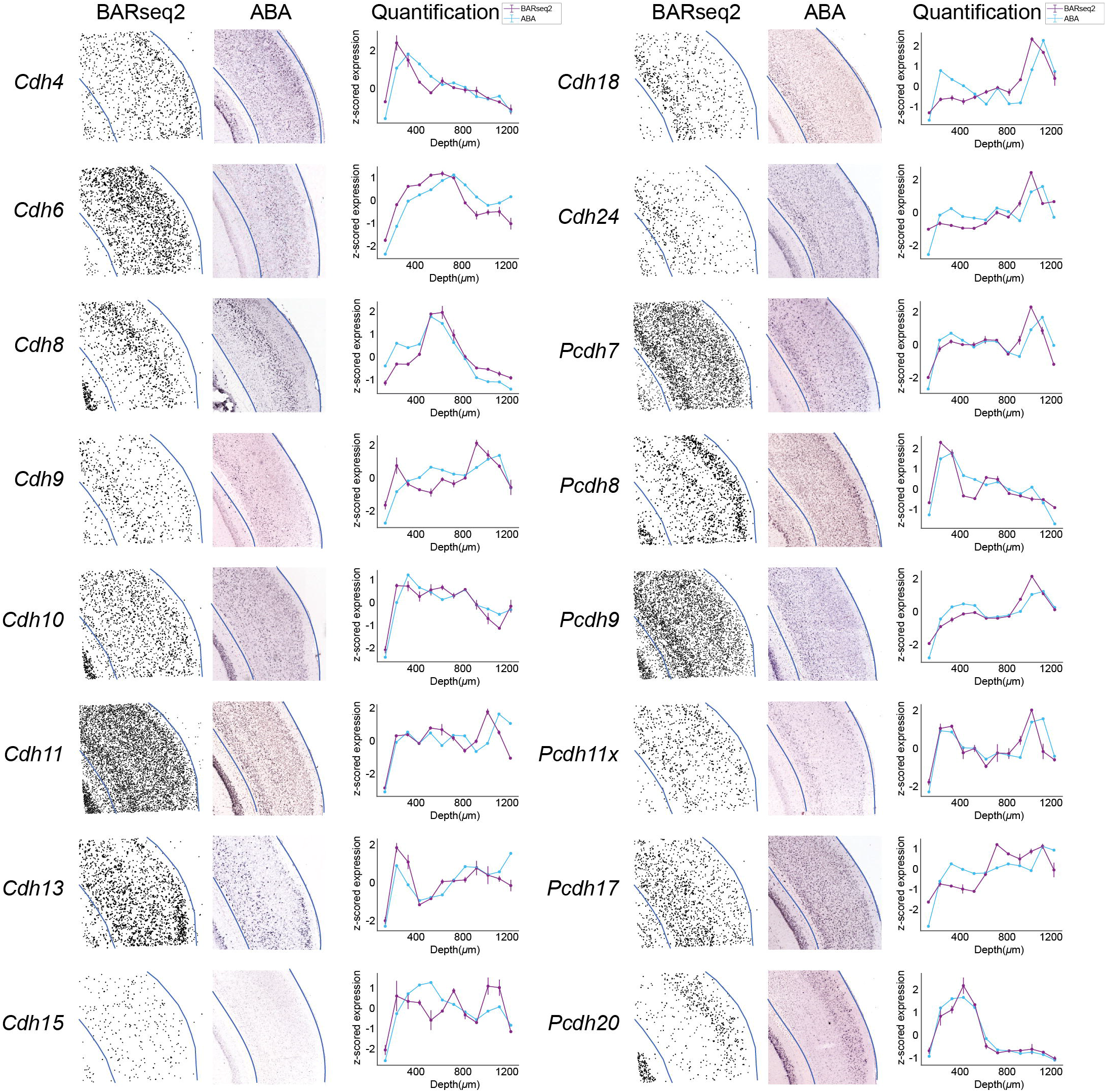
Comparison between BARseq2 and Allen gene expression atlas. Gene expression patterns in auditory cortex identified by BARseq2 are plotted next to *in situ* hybridization images of the same genes in Allen gene expression atlas (ABA) and the quantified laminar distribution of the gene in both datasets. Only genes that had coronal images in the Allen gene expression atlas are shown. Blue lines indicate the boundaries of the cortex in both BARseq2 and ABA images. In the laminar distribution plots, plotted values are means ± standard errors across two BARseq2 samples (purple lines) and one or multiple ABA samples (blue lines). Genes that had only a single sample in ABA do not have error bars.

**Extended Data Figure 4.**
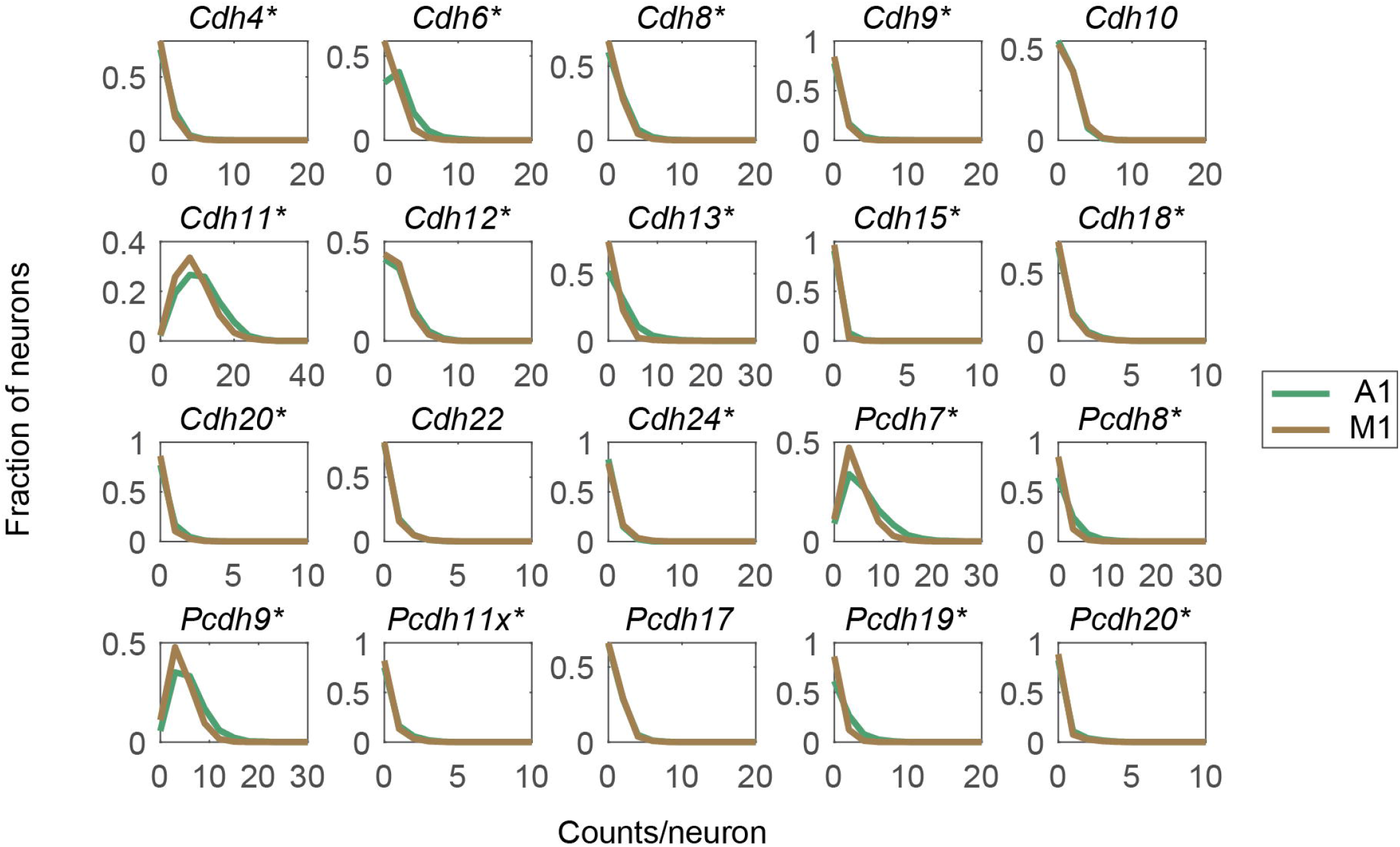
The distribution of read counts per cell for the indicated genes in auditory cortex (green) and motor cortex (brown). Asterisks indicate genes with significant difference in expression between the two areas (p < 0.05 using rank sum test after Bonferroni correction).

**Extended Data Figure 5.**
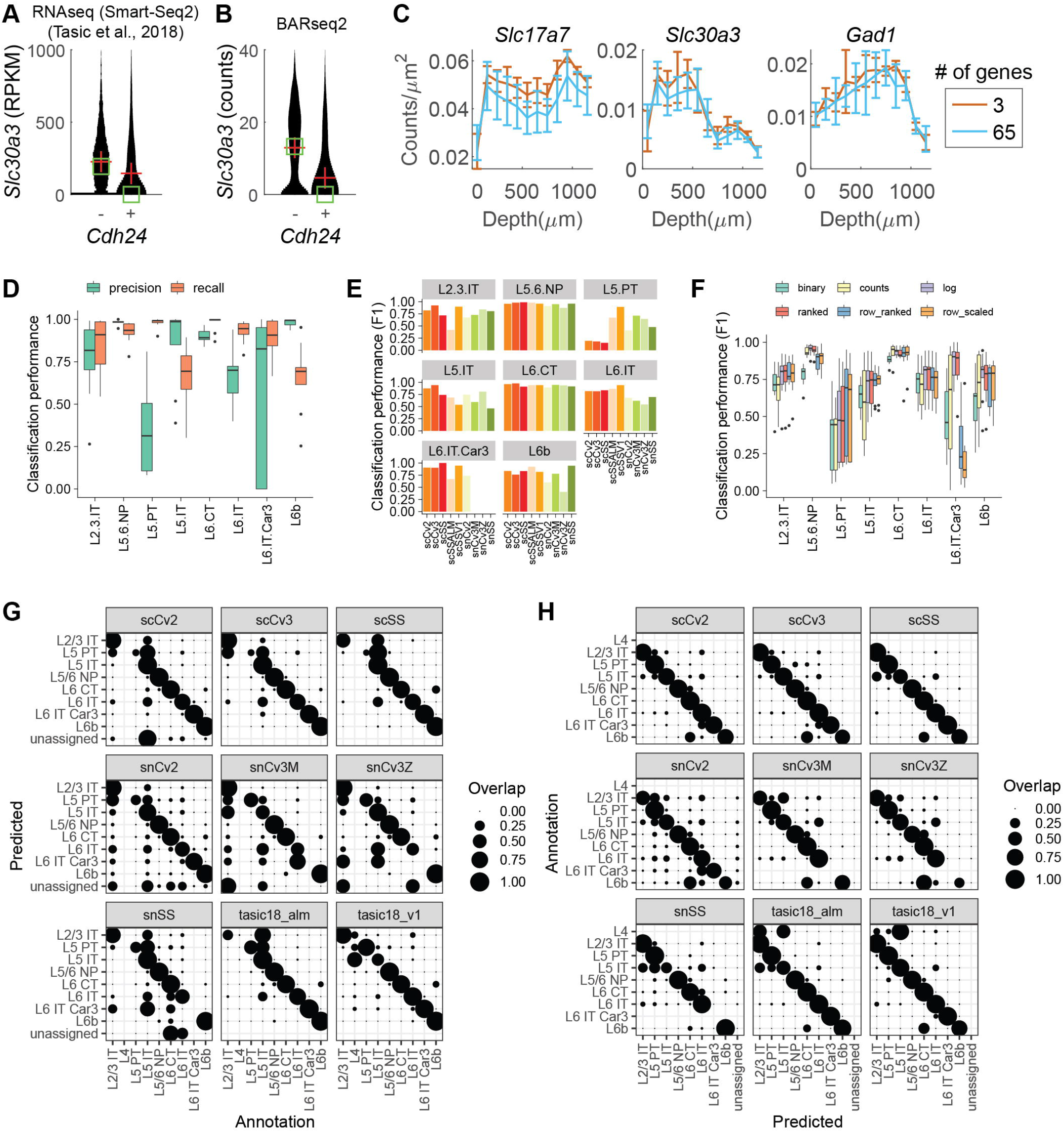
Transcriptomic typing using BARseq2. (A)(B) *Slc30a3* expression in excitatory neurons with or without *Cdh24* expression in single-cell RNAseq (A) from Tasic et al. (2018) or in BARseq2 (B). A cell is considered expressing *Cdh24* if the expression is higher than 10 RPKM in RNAseq or 1 count in BARseq2. Red crosses indicate means and green squares indicate medians. (C) Expression density (mean ± standard deviation) across laminar positions for the indicated genes. n = 3 slices for the three-gene panel and n = 5 slices for the 65-gene panel. (D) Precision and recall of cell typing using the marker gene panel across nine single cell datasets. (E) Breakdown of average performance for each cell type in each dataset. The datasets are: scSSALM and scSSV1 are single cell SmartSeq datasets from ALM and V1 respectively (Tasic et al., 2018). All other datasets are BICCN M1 datasets (Yao et al., 2020) and the name indicates the technology used (sc = single cell, sn = single nuclei, Cv2/3 = Chromium v2/3, SS = SmartSeq). (F) Average cell typing performance for six normalization strategies. (G) Confusion matrix showing overlap between prediction and annotations, normalized by predictions. This plot emphasizes precision; it indicates the probability that a given prediction was correct. (H) Confusion matrix showing overlap between prediction and annotations, normalized by annotations. This plot emphasizes recall; it indicates the probability that a given annotation was recovered.

**Extended Data Figure 6.**
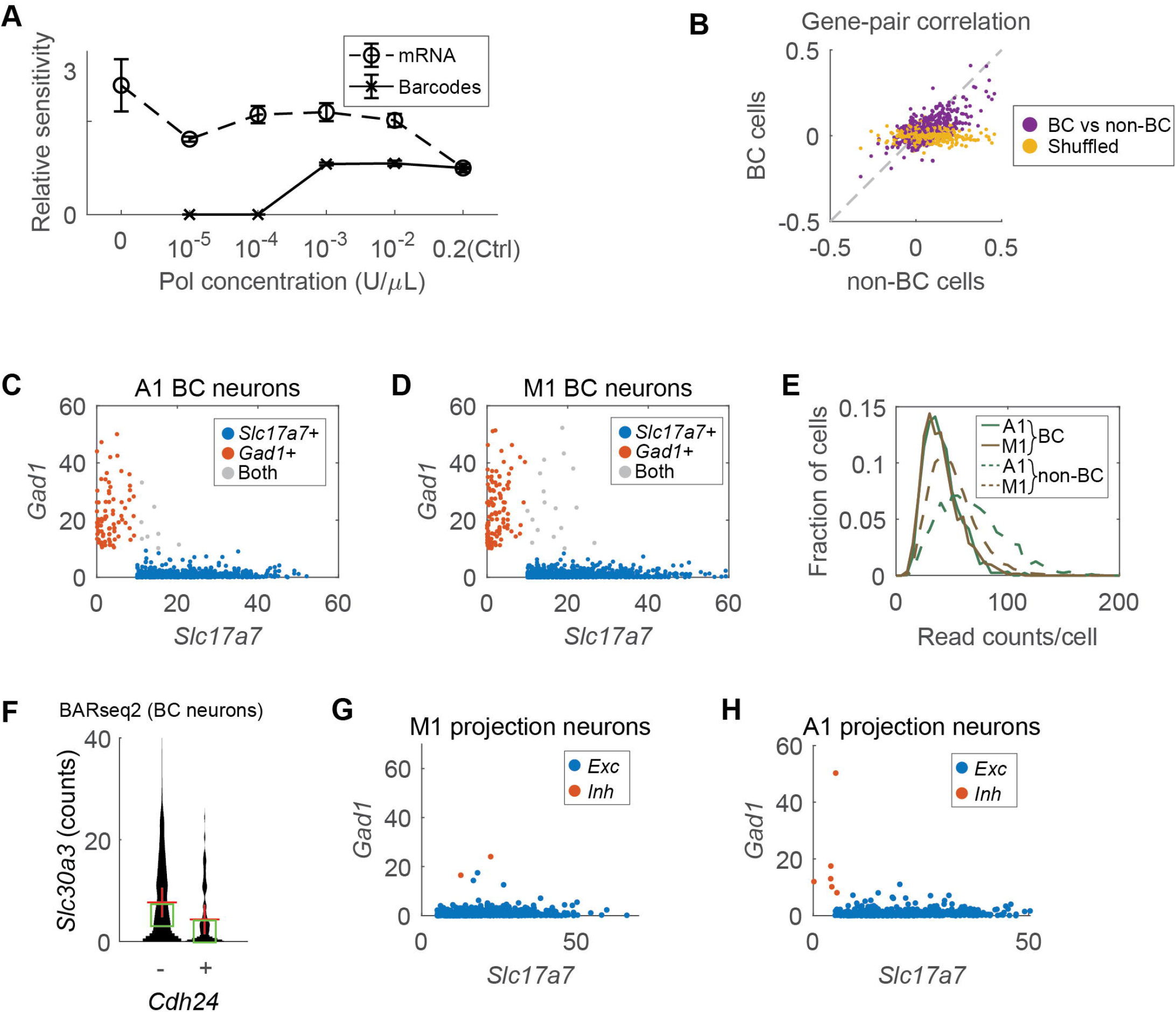
Correlating gene expression to projections using BARseq2. (A) Relative sensitivity of BARseq2 to barcodes (solid line) and endogenous mRNAs (dashed line) using the indicated concentration of Phusion DNA polymerase. Sensitivities are normalized to the original BARseq condition (*Ctrl*). Error bars indicate standard errors. n = 2 slices for each data point. (B) Correlation between pairs of genes in barcoded cells (y-axis) and in non-barcoded cells (x-axis) as determined by BARseq2. Shuffled data (yellow) are also plotted for comparison. (C)(D) *Slc17a7* (x-axes) and *Gad1* (y-axes) expression in barcoded neurons in auditory (C) or motor cortex (D). Only neurons with more than 10 counts in either gene are shown. (E) The distributions of read counts per barcoded neuron (solid lines) or non-barcoded neuron (dashed lines) in auditory (green) and motor (brown) cortex. (F) *Slc30a3* expression in barcoded excitatory neurons with or without *Cdh24* expression in BARseq2. A cell is considered expressing *Cdh24* if the expression is higher than 1 count. Red crosses indicate means and green squares indicate median. (G)(H) *Slc17a7* (x-axes) and *Gad1* (y-axes) expression in barcoded projection neurons in motor (G) or auditory cortex (H). Excitatory and inhibitory neurons are color-coded as indicated.

**Extended Data Figure 7.**
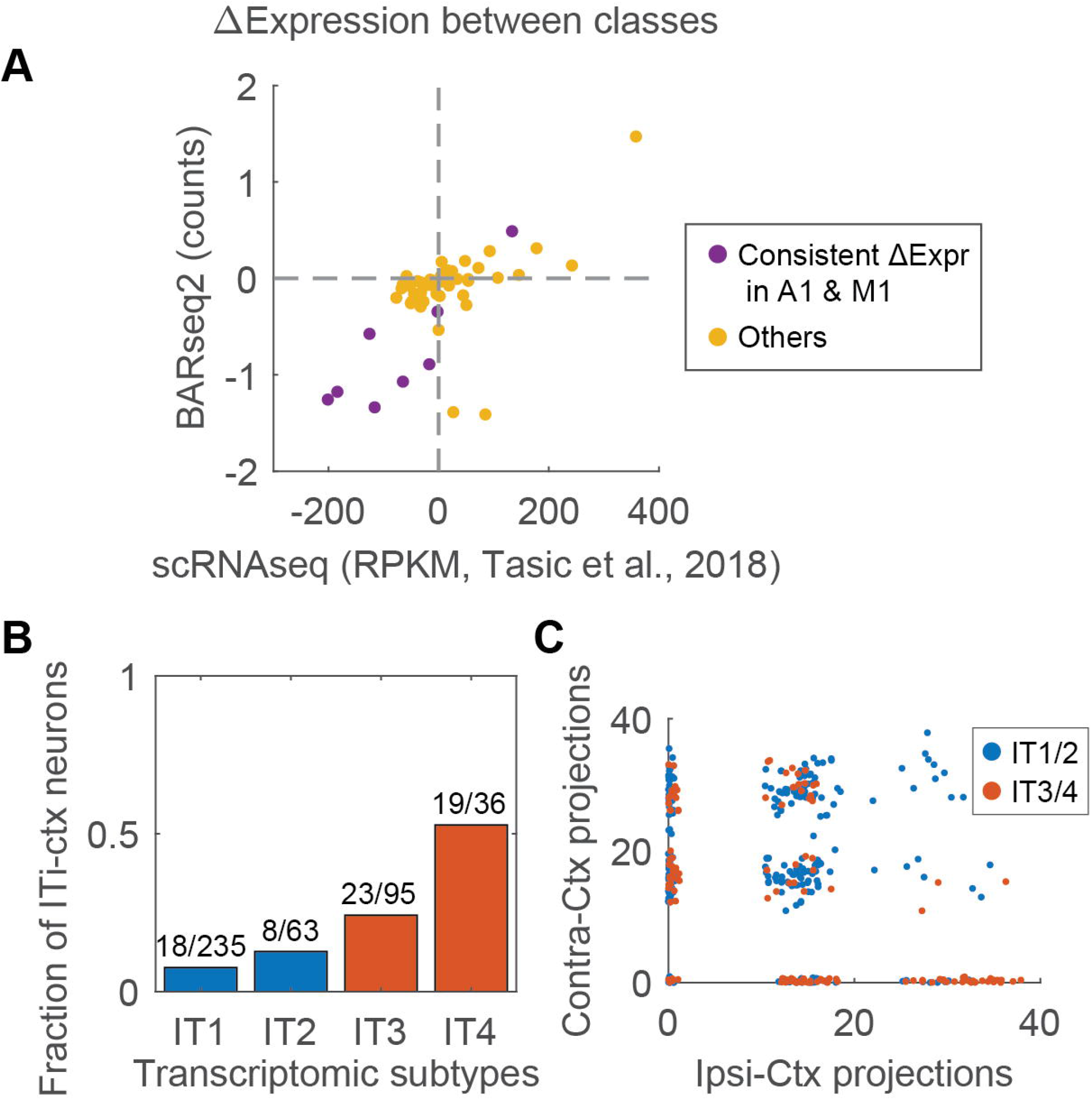
BARseq2 reveals projection and gene expression differences across major classes and IT subtypes. (A) Differential gene expression across major classes (IT, PT, and CT) observed using BARseq2 and single-cell RNAseq. Each dot shows the difference in mean expression of a gene across a pair of major classes observed using BARseq2 (y-axis) or single-cell RNAseq (x-axis). Differences in expression that were statistically significant (FDR < 0.05) in both A1 and M1 as shown by BARseq2 were labeled purple; otherwise they were labeled yellow. The single-cell RNAseq data used were collected in the visual cortex and anterior-lateral motor cortex (Tasic et al., 2018). (B) The fraction of ITi-ctx neurons in four transcriptomic types of IT neurons in auditory cortex. ITi-ctx neurons have only ipsilateral cortical projections and no striatal projections or contralateral projections (Chen et al., 2019). The number of ITi-ctx neurons and the total number of neurons for each transcriptomic type are labeled on top of the bars. (C) The projection strengths for contralateral (y-axis) and ipsilateral (x-axis) cortical projections for each IT neuron in auditory cortex. IT1/IT2 neurons are labeled blue and IT3/IT4 neurons are labeled red.

**Extended Data Figure 8.**
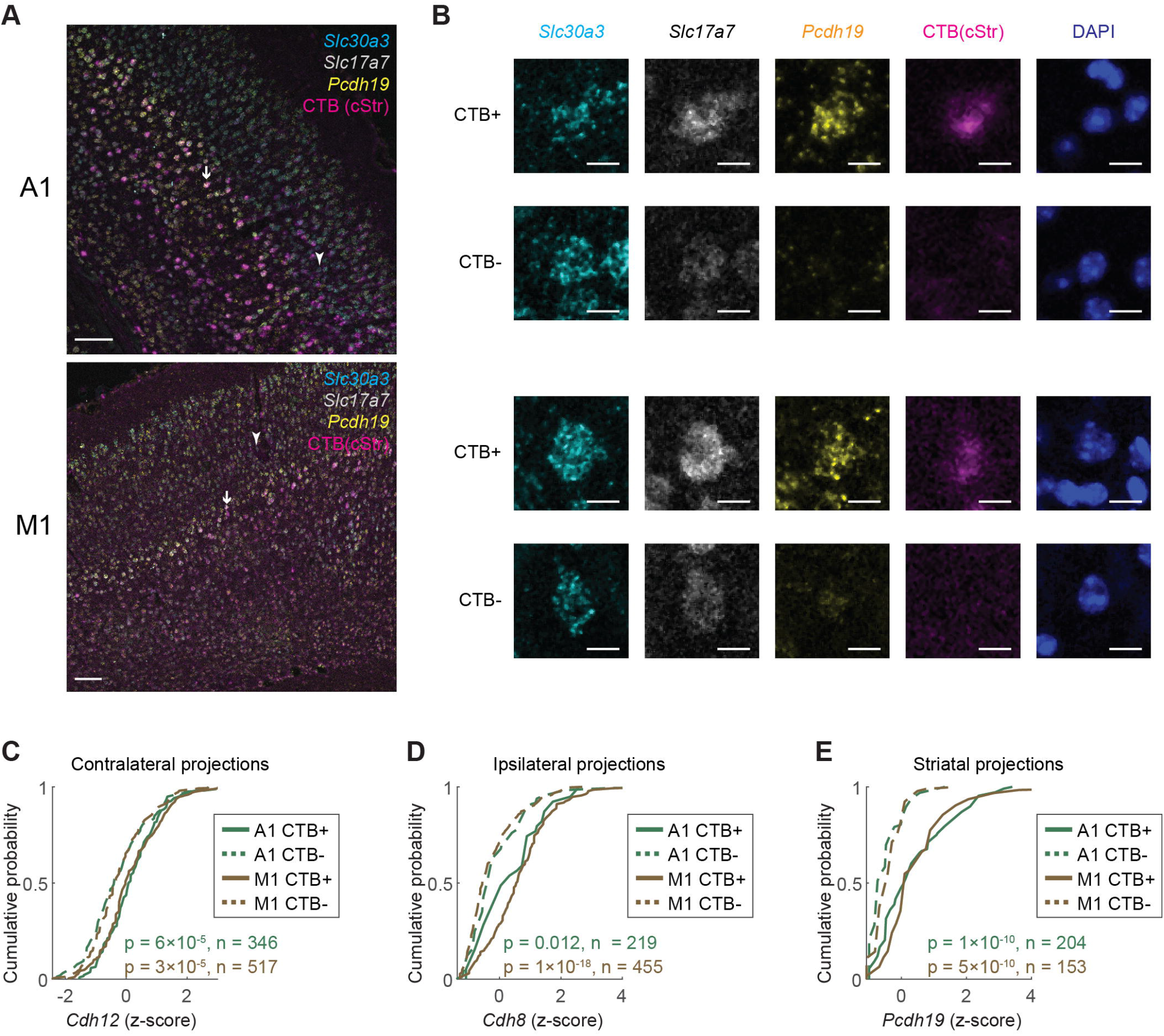
Validation of correlation between cadherins and IT projections. (A) Representative images of *in situ* hybridization in A1 (*top*) and M1 (*bottom*) slices with CTB labeling in the caudal striatum. Three marker genes and CTB labeling are shown in the indicated colors. Scale bars = 100 μm. Arrows and arrowheads indicate example CTB+ and CTB-neurons, respectively. (B) Crops of the indicated individual channels of example neurons from (A). Scale bars = 10 μm. (C)-(E) Cumulative probability distribution of the expression of *Cdh12* (C), *Cdh8* (D), and *Pcdh19* (E) in neurons with or without retrograde labeling of contralateral (C), ipsilateral (D), or caudal striatal (E) projections. p values from rank sum tests after Bonferroni correction and numbers of neurons used for each experiment are indicated. N = 2 animals for each experiment.

**Extended Data Figure 9.**
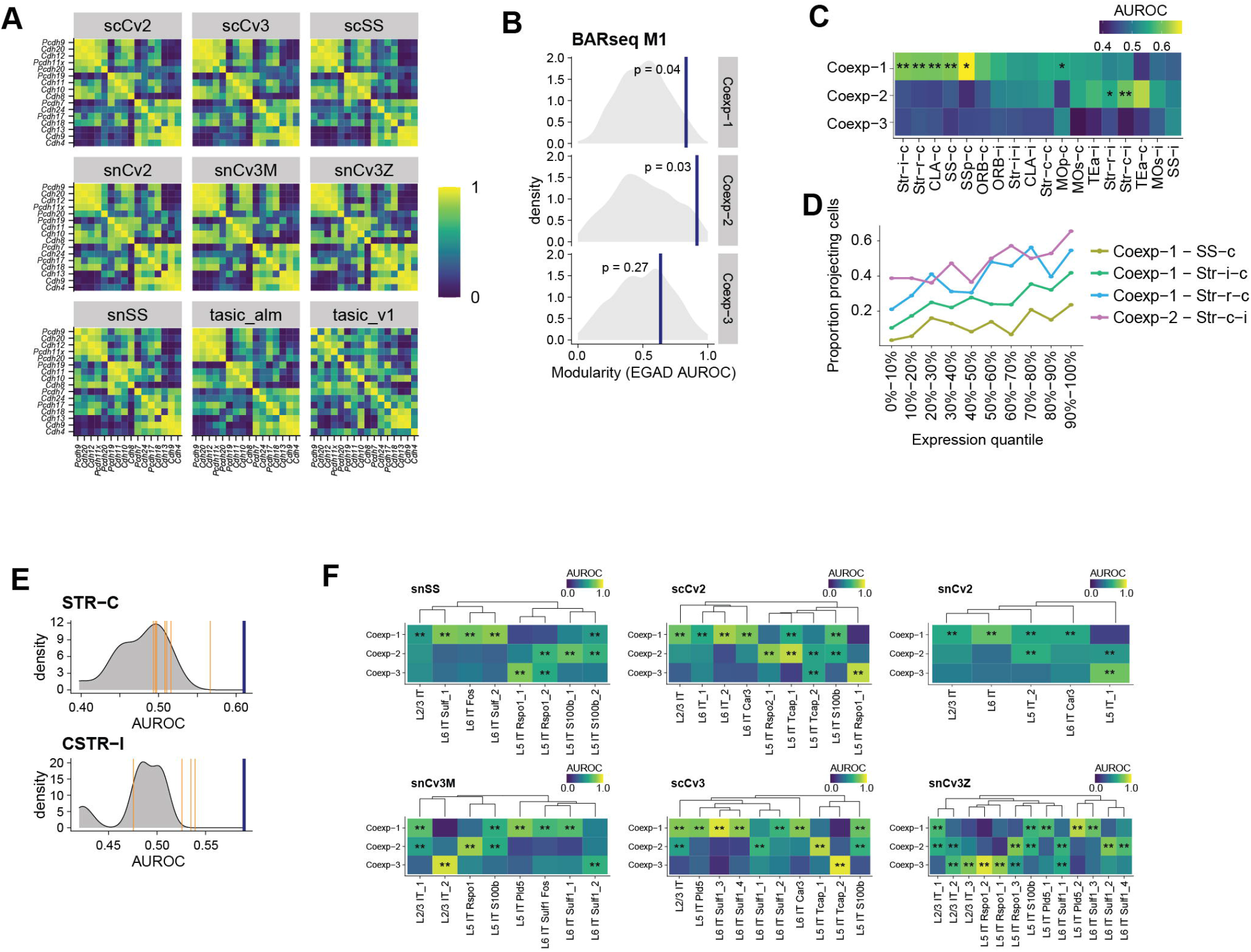
Cadherin co-expression modules correlate with IT projections. (A) Correlation among cadherins in IT neurons in motor cortex identified in the indicated single-cell RNAseq datasets (Tasic et al., 2018; Yao et al., 2020). The datasets included are: tasic_alm and tasic_v1 are single cell SmartSeq datasets from ALM and V1 respectively (Tasic et al., 2018); all other datasets are BICCN M1 datasets (Yao et al., 2020); the name indicates the technology used (sc = single cell, sn = single nuclei, Cv2/3 = Chromium v2/3, SS = SmartSeq). (B) Modularity (EGAD AUROC) of co-expression modules in BARseq2 M1 against null distribution of modularity (node permutation). BARseq2 modularity is shown by the blue lines with the corresponding p-values. (C) Association (AUROC) between cadherin co-expression modules and the indicated projections. Significant associations are marked by asterisks (* FDR < 0.1, ** FDR < 0.05). (D) Fractions of neurons with the indicated projections as a function of co-expression module expression. (E) Distribution of associations of the indicated projection modules with gene expression. Association with significant gene module is shown by a blue line; association with single genes from that module is shown by orange lines; association with all other genes is shown by a gray density. (F) Association of the three co-expression modules in transcriptomic IT neurons in the indicated datasets (AUROC, significance shown as in C).

## Methods

### Animal processing and tissue preparation

All animal procedures were carried out in accordance with the Institutional Animal Care and Use Committee protocol 19-16-10-07-03-00-4 at Cold Spring Harbor Laboratory. A list of animals used is provided in Supp. Table S1.

For samples used for only endogenous mRNA detection, 8-10 week old male C57BL/6 mice were anesthetized and decapitated. We immediately embedded the brain in OCT in a 22 mm^2^ cryomold and snap-froze the tissue in an isopentane bath submerged in liquid nitrogen. Sections were cut into 10 μm-thick slices on Superfrost Plus Gold Slides (Electron Microscopy Sciences). Unlike in the original BARseq, the sections were directly melted onto slides without the use of a tape transfer system. This change in mounting methods allowed increased efficiency in gene detection. The slides were stored at −80 °C until use.

For BARseq2 samples, 8-10 week old male C57BL/6 mice were injected in the left auditory cortex at −2.6 mm AP, −4.3 mm ML from the bregma, with 140 nL 1:3 diluted Sindbis virus at depths 300 μm, 500 μm, and 800 μm at a 30 ° angle. After 24 hrs, we anesthetized and decapitated the animal, punched out the injection site, and snap-froze the rest of the brain on a razor blade on dry ice for conventional MAPseq (Chen et al., 2019). The injection site was embedded, cryo-sectioned, and stored as described above.

To prepare samples for BARseq2 experiments, we immersed slides from −80 °C instantly into freshly made 4 % PFA (10mL vials of 20 % PFA; Electron Microscopy Sciences) in PBS for 30 mins at room temperature. We washed the samples in PBS for 5 mins before installing HybriWell-FL chambers (22 mm × 22 mm × 0.25 mm; Grace Bio-labs) for subsequent reactions on the samples. We then dehydrated the samples in 70 %, 85 %, and 100 % EtOH for 5 mins each, and then washed in 100 % EtOH for at least 1 hr at 4 °C. Finally, we rehydrated the samples in PBST (0.5 % Tween-20 in PBS).

For retrograde labeling experiments, we prepared 1.0 mg/mL of Cholera Toxin subunit B (CTB) in PBS from 100 μg for injections (see Supp. Table S3 for a list of animals and coordinates used). We perfused the animals with fresh 4 % PFA 96 hrs after injection, post-fixed for 24 hrs in 4 % PFA, and cryo-protected in 10 % sucrose in PBS for 12 hrs, 20 % sucrose in PBS for 12 hrs, and 30 % sucrose in PBS for 12 hrs. The brain was then frozen in OCT and cryo-sectioned to 20 μm slices using a tape transfer system.

### BARseq2 detection of endogenous genes

We prepared a master mix of reverse transcription primers at 0.5 μM each for all target mRNAs. For volumes exceeding the amount required for reverse transcription, we speed-vacuumed to concentrate the primer mix into a smaller volume. We then prepared the reaction [0.5 μM per gene RT primer (IDT), 1 U/μL RiboLock RNase inhibitor (Thermo Fisher Scientific), 0.2 μg/μL BSA, 500 μM dNTPs (Thermo Fisher Scientific), 20 U/μL RevertAid H-Minus M-MuLV reverse transcriptase (Thermo Fisher Scientific) in 1× RT buffer]. We incubated the samples in reverse transcription at 37 °C overnight. After reverse transcription, we crosslinked the cDNAs in 50 mM BS(PEG)_9_ (Thermo Fisher Scientific) for 1 hr and neutralized excess crosslinker with 1 M Tris-HCl, pH 8.0 for 30 mins, and then washed the sample with PBST twice to eliminate excess Tris. We then prepared a master padlock mix with 200 nM per padlock probe for each target mRNA and speed-vacuumed the mixture for a higher concentration at a smaller volume, if necessary. We ligated the gene padlock probes on the cDNA [200 nM per gene padlock (IDT), 1 U/μL RiboLock RNase Inhibitor, 20 % formamide (Thermo Fisher Scientific), 50 mM KCl, 0.4 U/μL RNase H (Qiagen), and 0.5 U/μL Ampligase (Epicentre) in 1× Ampligase buffer] for 30 mins at 37 °C and 45 mins at 45 °C. Finally, we performed rolling circle amplification (RCA) [125 μM amino-allyl dUTP (Thermo Fisher Scientific), 0.2 μg/μL BSA, 250 μM dNTPs, 5 % glycerol, and 1 U/μL ◻29 DNA polymerase (Thermo Fisher Scientific) in 1× ◻29 DNA polymerase buffer] overnight at room temperature. After RCA, we again crosslinked the rolonies in 50 mM BS(PEG)_9_ for 1 hr, neutralized with 1 M Tris-HCl, pH 8.0 for 30 mins, and washed with PBST. We washed the sample in hybridization buffer [10 % formamide in 2× SSC] and then either added probe detection hybridization solution [0.25 μM fluorescent probe in hybridization buffer] or genes sequencing primer hybridization solution [1 μM of sequencing primer in hybridization buffer] for 10 mins at room temperature. We then washed the sample with hybridization buffer three times at two mins each, rinsed the sample in PBST twice, and proceeded to imaging or continue with Illumina sequencing.

### BARseq2 simultaneous detection of endogenous genes and barcodes

We prepared a master mix of reverse transcription primers at 0.5 μM each for all target mRNAs. For volumes exceeding the amount required for reverse transcription, we speed-vacuumed to concentrate the primer mix into a smaller volume. We then prepared the reaction [0.5 μM per gene RT primer (IDT), 1 μM barcode LNA RT primer (Qiagen), 1U/μL RiboLock RNase inhibitor (Thermo Fisher Scientific), 0.2 μg/μL BSA, 500 μM dNTPs (Thermo Fisher Scientific), 20 U/μL RevertAid H-Minus M-MuLV reverse transcriptase (Thermo Fisher Scientific) in 1× RT buffer], adding the barcode LNA primer last into the reaction mix to reduce cross-hybridization due to the LNA strong binding affinity. We incubated the samples in reverse transcription at 37 °C overnight. After reverse transcription, we crosslinked the cDNAs in 50 mM BS(PEG)_9_ (Thermo Fisher Scientific) for 1 hr and neutralized excess crosslinker with 1 M Tris-HCl, pH 8.0 for 30 mins, and then washed the sample with PBST twice to eliminate excess Tris. We then prepared a master padlock mix with 200 nM per padlock probe for each target mRNA and speed-vacuumed the mixture for a higher concentration at a smaller volume, if necessary. We ligated the gene padlock probes on the cDNA [200 nM per gene padlock (IDT), 1 U/μL RiboLock RNase Inhibitor, 20 % formamide (Thermo Fisher Scientific), 50 mM KCl, 0.4 U/μL RNase H (Qiagen), and 0.5 U/μL Ampligase (Epicentre) in 1× Ampligase buffer] for 30 mins at 37 °C and 45 mins at 45 °C. After ligating padlock probes for our target genes, we ligated the padlock probe for the barcode cDNA [100 nM barcode padlock (IDT), 50 μM dNTPs, 5 % glycerol, 1 U/μL RiboLock RNase Inhibitor, 20 % formamide (Thermo Fisher Scientific), 50 mM KCl, 0.4 U/μL RNase H (Qiagen), 0.001 U/μl Phusion DNA polymerase (NEB), and 0.5 U/μL Ampligase (Epicentre) in 1× Ampligase buffer] without any wash in between, and incubated the reaction for 5 mins at 37 °C and 40 mins at 45 °C. We then washed the sample twice with PBST and once with hybridization buffer [10 % formamide in 2× SSC], before hybridizing 1 μM of RCA primer in hybridization buffer for 15 mins at room temperature. We washed the sample with hybridization buffer three times at two mins each. Finally, we performed rolling circle amplification (RCA) [125 μM aadUTP (Thermo Fisher Scientific), 0.2 μg/μL BSA, 250 μM dNTPs, 5 % glycerol, and 1 U/μL D29 DNA polymerase (Thermo Fisher Scientific) in 1× D29 DNA polymerase buffer] overnight at room temperature. After RCA, we again crosslinked the rolonies in 50 mM BS(PEG)_9_ for 1 hr, neutralized with 1 M Tris-HCl, pH 8.0 for 30 mins, and washed with PBST. We washed the sample in hybridization buffer [10 % formamide in 2× SSC] and then added genes sequencing primer hybridization solution [1 μM of sequencing primer in hybridization buffer] for 10 mins at room temperature. We then washed the sample with hybridization buffer three times at two mins each, rinsed the sample in PBST twice, and proceeded to Illumina sequencing.

### *In situ* sequencing of endogenous genes

To sequence the endogenous genes using Illumina sequencing chemistry, we used the HiSeq Rapid SBS Kit v2 reagents to reduce cost from the original sequencing protocol (Chen et al., 2019). For the first cycle, we incubated samples in Universal Sequencing Buffer (USB) at 60 °C for 3 mins, then washed in PBST, and then incubated in iodoacetamide (9.3 mg in 2 mL PBST) at 60 °C for 3 mins. We washed the sample in PBST again, rinsed with USB twice more, and then incubated in Incorporation Mix (IRM) at 60 °C for 3 mins. We repeated the IRM step again to ensure as close to 100 % complete reaction as possible. We then washed the sample in PBST once and then continued to wash in PBST four more times at 60 °C for 3 mins each time. To reduce bleaching during imaging, we imaged the sample in Universal Scan Mix (USM).

For subsequent cycles, we first washed samples in USB, then incubated in Cleavage Reagent Mastermix (CRM) at 60 °C for 3 mins. We repeated the CRM step to ensure complete reaction and washed out residual CRM twice with Cleavage Wash Mix (CWM). We then washed the sample with USB, and then with PBST, before incubating in iodoacetamide at 60 °C for 3 mins. We repeated this step again to ensure we block as many of the free thiol-groups as possible to reduce background. We then continued with IRM and PBST washes as described for the first cycle and imaged after each cycle. We performed four sequencing cycles and seven sequencing cycles in total for our cadherins panel of 23 genes and our motor cell type markers and cadherins panel of 65 genes, respectively.

To visualize high expressors, we cleaved the fluorophores in the fifth sequencing cycle and washed the sample with CWM and PBST. We then washed our sample in hybridization buffer and added probe detection solution (0.5 μM each probe in hybridization buffer) for four different fluorescent probes detecting *Slc17a7*, *Gad1*, *Slc30a3*, and all previously sequenced genes, respectively, for 10 mins at room temperature. We washed the sample in the same hybridization buffer three times for two mins each, washed in PBST, before adding DAPI stain (ACDBio) for 2 mins at room temperature. We rinsed in PBST again and finally in USM for imaging.

### *In situ* sequencing of barcodes

After sequencing and hybridizing for endogenous genes as described above, we stripped the sample of all hybridized oligos and sequenced bases by incubating twice in strip buffer (40 % formamide in 2× SSC with 0.01 % Triton-X) at 60 °C for 10 mins. We washed with PBST, then washed with hybridization buffer, and then incubated samples in barcode sequencing primer hybridization solution (1 μM sequencing primer in hybridization buffer) for 10 mins at room temperature. We washed with hybridization solution three times for two mins each, before rinsing twice in PBST. We sequenced barcodes with the same sequencing procedure as described for endogenous genes but for 15 cycles in total. Around cycle 4 or 5, we eliminate the iodoacetamide blocker incubation for the rest of sequencing because iodoacetamide blockage is irreversible, so further incubation in this blocker becomes unnecessary after several cycles.

### Target area barcode sequencing

Barcode sequencing in target brain areas were sequenced by the Cold Spring Harbor Laboratory MAPseq core following procedures used in a previous study (Chen et al., 2019). The target areas were dissected to match two other studies in A1 (Chen et al., 2019) and in M1 (Chen et al., unpublished observations). Detailed description of each dissected area and correspondence to the Allen reference atlas are shown in Supp. Table S2. Example annotated images from the dissected brain slices are provided at Mendeley Data (see Data Availability).

### Fluorescent *in situ* hybridization (FISH)

FISH experiments were performed using RNAscope Fluorescent Multiplex Kit v1 according to the manufacturer’s protocols with minor modifications to sample preprocessing. For FISH experiments in comparison to BARseq2 endogenous mRNA detection (Fig. 1F; Fig. 2E), the samples were fresh-frozen in isopentane bath as described above. From −80 °C storage, the samples were immediately submerged in freshly-made 4 % PFA (Electron Microscopy Sciences) for 15 mins at 4 °C, then dehydrated in 75 %, 85 %, and 100 % ethanol twice for 5 mins each. After air-drying, we assembled HybriWell-FL chambers (22 mm × 22 mm × 0.25 mm; Grace Bio-Labs) and digested the samples in Protease IV for 30 mins at room temperature. We washed the samples in PBST, and then proceeded with probe hybridization and subsequent amplification and visualization steps following the manufacturer’s protocol, and mounted the samples with coverslips finally for imaging.

For FISH experiments in retrogradely labeled samples, we first imaged the samples before performing FISH. The samples were then dehydrated in 50 %, 75 % and 100 % ethanol twice for 5 mins each. After air-drying the samples, we either assembled HybriWell-FL chambers (22 mm × 22 mm × 0.25 mm; Grace Bio-Labs) or drew a barrier around the samples using a ImmEdge hydrophobic barrier pen. The samples were then digested in Protease III for 30 mins at 40 °C, and washed in nuclease-free H_2_O twice. We then proceeded to probe hybridization and subsequent amplification and visualization steps following the manufacturer’s protocol, and mounted the samples with coverslips finally for imaging.

For Fig. 1F, the FISH probes used were Mm-Slc17a7-C1, Mm-Slc30a3-C2, and Mm-Cdh13-C3 visualized with Amp4 A It A. For Fig. 2E, the FISH probes used were Mm-Pcdh19-C1, Mm-Cdh8-C2, and Mm-Pcdh20-C3 visualized with Amp4 A It A. For retrograde labeling experiments in Fig. S9A-E, the FISH probes used for the cadherins were Mm-Cdh12-C1 (custom-ordered no. 842531), Mm-Cdh8-C1, or Mm-Pcdh19-C1, in addition to Mm-Slc30a3-C2 and Mm-Slc17a7-C3, visualized with Amp4 A It C.

### Imaging

All sequencing experiments were performed on a Olympus IX81 microscope with Crest X-light 2 spinning disk confocal, a Photometrics BSI prime camera and an 89North LDI 7-channel laser bank. Retrograde labeling experiments were imaged either on the same microscope or on an LSM 710 Laser Scanning confocal microscope. Filters and lasers used for imaging are listed in Supp. Table S4.

For all BARseq2 experiments, we imaged endogenous genes using an Olympus UPLFLN 40× 0.75 NA air objective and tiled 5 × 5 with 15 % overlap between tiles for all sequencing cycles and the hybridization cycles. For each sequencing cycle, the four sequencing channels (G, T, A, and C) and the DIC channel was captured. For hybridization cycles, GFP, RFP, TexasRed, Cy5, and DIC channels were captured. At the last cycle (usually the hybridization cycle for high expressors), we also imaged the DAPI channel.

For barcode sequencing, we imaged the first three cycles using the same imaging settings described above at 40×. The third sequencing cycle was additionally reimaged at 10× using an Olympus UPLANAPO 10× 0.45 NA air objective without tiling. All subsequent barcode sequencing cycles were all imaged at 10×.

On the spinning disk confocal, all 40× BARseq2 and FISH images were acquired as z-stacks with 1 μm step size and 0.16 μm xy pixel size, and all 10× images were acquired as z-stacks with 5 μm step size.

On the LSM 710, CTB labeled samples were first imaged using a Plan-Apochromat 10× 0.45 NA objective without a coverslip as a z-stack with 7 μm z-step size and 0.7 μm xy pixel size. After FISH, the same samples were imaged using a Plan-Apochromat 20× 0.8 NA objective as a z-stack with 2 μm step size and 0.35 μm xy pixel size.

### Probe design

We designed reverse transcription primers and padlock probes for sequencing barcodes as described previously (Chen et al., 2019) (see Supp. Table S1).

To design reverse transcription primers and padlock probes, we tried to design as many probe sets as possible on each transcript while avoiding the end (~20 nt) of the mRNA transcripts and ensuring at least a 3 nt gap between two adjacent probe sets. Specific reverse transcription primers were designed to be 25 to 26 nt with amino modifier C6 at the 5’ end and HPLC purified. In addition, we avoided sequences that contained G/C quadruplexes and/or had a low melting temperature (below 55 °C). Padlock probes were designed to have two arms of 21 to 23 nt with minimum T_m_ of 58 °C, GC contents between 40 % and 60 %, and high complexity. The two arms were connected by a backbone consisting of a 32 nt sequencing primer or detection probe target site, a 7 nt gene-specific index, and a 3 nt 3’ linker. For padlock probes designed for hybridization readout, different backbone sequences were used for different genes. We further filtered out padlock probe sequences with potential non-specific binding. To find potential non-specific binding targets, we blasted the ligated padlock arm sequences against the mouse genome and identified all targets with (1) 3 nt of perfect match on either side of the ligation junction, (2) no gap and/or insertion within 7 nt of the ligation junction, and (3) melting temperatures of at least 37 °C for non-specific binding of each arm.

The gene-specific indices were chosen so that all cadherin genes have a minimum Hamming distance of two by sequencing the first four bases, and all cadherins and cell-type marker genes have a minimum Hamming distance of two by sequencing the first five bases and three by all seven bases.

We maximized the number of padlock probe sets for *Slc17a7* (23 probes), *Slc30a3* (19 probes), *Gad1* (24 probes), and *Cdh13* (30 probes). These probe sets were used to evaluate the relationship between detection sensitivity and probe numbers. For the cadherin panels and the cell-type marker panels, we selected a subset of probes for each gene so that we have at most 12 probe sets per gene. Some shorter genes had fewer than 12 probes.

A detailed description of probe sets used for each experiment and their sequences is provided in Supp. Table S1.

### Single-cell RNAseq of auditory cortex

Single-cell RNAseq experiments were performed as described previously (Chen et al., 2019) using 10x Genomics Chromium Single Cell 3′ Kits v3. One of the single-cell RNAseq dataset was previously published (Chen et al., 2019), and a new dataset was obtained in this study.

### BARseq2 data processing

MAPseq data were processed as described previously (Chen et al., 2019; Kebschull et al., 2016). A sample script for processing MAPseq datasets is provided at Mendeley Data (see Data Availability).

To process *in situ* sequencing data for genes, we first performed max projection of the image stacks along the z-axis. Each max projection image was then corrected for sequencing channel bleedthrough and lateral shift across channels. The images were then filtered with a median filter and background subtracted using a rolling ball with a radius of 10 pixels. The sequencing cycle images were then registered to the first sequencing cycle using the sum of all four sequencing channels, and the hybridization images were registered to the first sequencing using the channel that labeled all sequenced rolonies. Registrations were performed by maximizing enhanced cross correlation (Evangelidis and Psarakis, 2008). After all images were registered, putative rolonies were then picked from the first sequencing cycle by finding all peaks that were at least brighter than all surrounding valleys by a certain threshold determined empirically. This was achieved by first performing morphological reconstruction using the original image as mask and the image minus the threshold as marker, followed by identifying all local maxima. We then deconvolve all registered images and find the signal intensities for all rolonies across all sequencing cycles and channels.

At this point the signal for each rolony is represented by an m × 1 vector, in which m equals four (sequencing channels) times the number of cycles. To identify the gene that each rolony correspond to, we project the signal vector onto the signal vector of all genes and find the two genes with the highest projections, I1 and I2. For rolonies whose (I1 - I2) / I1 is above a threshold, we assign the genes with the highest projections to these rolonies. The remaining rolonies are filtered out. For hybridization cycles, the channel in which the rolonies are found is used directly to identify the genes.

In experiments in which genes were detected without barcodes for projection mapping, we segmented cell bodies using both the DAPI signals and the sequencing signals with Cellpose (Stringer et al., 2020).

In experiments in which genes were detected in conjugation with barcodes, we further registered barcode sequencing cycles to the first sequencing cycle for genes using the DIC channel. The barcode sequencing images were then filtered with a median filter and background subtracted using a rolling ball with a radius of 50 pixels. The high-resolution images for the second and third cycles were then registered to the first sequencing cycle of barcodes using the sum of all four sequencing channels. The low-resolution images of the third sequencing cycle were then registered to the high-resolution image of the same cycle.

To segment the barcoded cells from the high-resolution images, we first identify “seed” pixels by identifying local maxima in the first sequencing cycle image as described above. These seed pixels are positions of the strongest signal within putative cell bodies. Then for each seed pixel, we calculate the projection of signal vectors for all other pixels within a local area on the signal vector of the seed pixel and the rejection of signal vectors for these pixels from the signal vector of the seed pixel. We then segment the cell bodies by finding all pixels that fulfill the following criteria: (1) the projections of their signal vectors are above a threshold, (2) the ratios between the rejections and projections are below a threshold, and (3) are connected to the seed pixel. In parallel, we perform a second segmentation using only the DAPI signals and gene sequencing images with marker-based watershedding without using the barcode sequencing images, and find the segmented cells that overlap with the barcode segmented cells. We then visually inspect the sequencing images and segmentations for each cell to determine which segmentation produced better result and to eliminate badly segmented cells. We then assign gene rolonies to the filtered segmented cells to produce the expression matrix.

To find the barcode sequences of the segmented cell, we integrate signals over the whole segmented cells and call the channel with the strongest signal as the base in both the high-resolution images and the low-resolution images. We then concatenate the sequences from the high-resolution images and the low-resolution images to produce the full barcode sequences. To find the projection patterns, these *in situ* sequenced barcodes are then matched to the barcodes identified in the projection areas allowing one mismatch but not ambiguous matches (i.e. one *in situ* barcode matching to multiple barcodes found in projection sites).

### Analysis of BARseq2 gene expression data

All analyses were carried out in MATLAB. Scripts for all analyses are provided at Mendeley Data (see Data Availability).

For analysis of gene-only datasets, neurons were first filtered by requiring at least 10 counts of *Slc17a7* or *Gad1* and were positioned within the cortex. To make the data comparable to previous studies (Chen et al., 2019), the cortical depths of neurons were normalized to a total thickness of 1200 μm for auditory cortex and 1500 μm for motor cortex.

To quantify the mutual exclusivity of *Slc17a7* and *Gad1* in neurons, we defined the exclusivity index E = P(Gad1|Slc17a7)/P(Gad1), where P(Gad1|Slc17a7) indicates the probability of a cell expressing at least 10 counts of *Gad1* conditioned on the expression of at least 10 counts of *Slc17a7*, and P(Gad1) indicates the probability of a cell expressing at least 10 counts of *Gad1* in all filtered neurons.

To find cadherins that were differentially expressed in cell types, the expression of cadherins in each cell type was compared to the expression of cadherins in all other cell types using rank sum tests.

To compare laminar distribution observed by BARseq2, FISH, and Allen Brain Atlas, we quantified gene expression signal densities across 100 μm bins in laminar depth. For BARseq2 and FISH, the quantification was done by counting dots. For Allen Brain Atlas, the quantifications were done by integrating signal intensities over all pixels in each bin. Because each bin had different number of pixels sampled in our data, we then divided the gene expression signals by the area observed in the images to calculate the density. We then z-scored the densities within each gene to produce the laminar profiles for each gene.

### Cell typing in BARseq2 and single cell data

To select a panel of marker genes, we chose meta-analytic markers from 7 single-cell RNAseq in the motor cortex (Yao et al., 2020), accessed from the NeMo archive as indicated in the manuscript. In each dataset and for each cell type, we extracted differentially expressed (DE) genes among excitatory neurons (“Glutamatergic” class, 1-vs-all DE, fold change > 2, Mann-Whitney FDR < 0.05). We filtered out lowly expressed genes (average Counts Per Million < 100), then ranked genes according primarily by the number of datasets where they were DE, secondarily by average fold change and selected the top 5 markers.

To call cell types in BARseq2 and single cell data, we used the following procedure. First we normalized counts to log(1 + CPM), then we computed the average marker expression for each cell type and assigned the cell type with the highest average expression. If two marker sets were tied for highest expression, the cell was left unassigned.

### Analysis of BARseq2 gene expression and projection dataset

For analysis of BARseq2 datasets with both gene expression and projections, we first evaluated the mutual exclusivity of *Slc17a7* and *Gad1* expression as stated above. For this purpose, the neurons were filtered with the same thresholds as in the gene-only dataset. For all other analyses, we used a more relaxed filtering to compensate for the reduced gene expression in barcoded cells, requiring neurons to have at least 5 counts of *Slc17a7* or *Gad1*. In this filtered set, neurons were considered excitatory if the counts of *Slc17a7* were larger than the counts of *Gad1*, and were considered inhibitory if the counts of *Gad1* were larger than the counts of *Slc17a7*. Projection data were log normalized as in previous studies (Chen et al., 2019). We further normalized the projection strengths of each area to two previous clustered BARseq dataset (Chen et al., 2019) and used a random forest classifier to assign neurons to projection clusters.

To identify projection differences across transcriptionally defined IT subtypes in auditory cortex, we used definitions for four IT subtypes that were consistent with a previous study (Chen et al., 2019) to allow easy comparison. Specifically, we defined IT1 as neurons with depths less than 590 μm, IT2 as neurons with depths between 590 and 830 μm and did not express *Cdh13*, IT3 as neurons between 590 and 830 μm that expressed *Cdh13* or neurons deeper than 830 μm that expressed *Slc30a3*, and IT4 as neurons deeper than 830 μm that did not express *Slc30a3*.

To find cadherins that were differentially expressed across major projection classes and between auditory and motor cortex, we performed rank sum tests for pairwise comparisons among major classes or the two areas for each cadherin and calculated the FDRs.

Projection modules were identified using non-negative matrix factorization (Lee and Seung, 1999). To find cadherins that were associated with projection modules, we calculated the Spearman correlation between the coefficients for projection modules and gene counts. To generate the plots of differential gene expression in Fig. 6E, we sorted the neurons by the coefficients for projection modules and smoothed gene expression using a window of 101 neurons.

To extract robust modules of co-expressed cadherins, we used a previously developed approach to combine multiple datasets meta-analytically, a crucial step to attenuate technical and biological noise (Ballouz et al., 2015; Crow et al., 2016). Briefly, we built co-expression networks using Spearman correlation for 7 single-cell RNAseq in the motor cortex (Yao et al., 2020), accessed from the NeMo archive as indicated in the manuscript and subset to the following subclasses: “L2/3 IT”, “L4/5IT”, “L5 IT”, “L6 IT” and “L6 IT Car3”. We ranked each network, then averaged the networks to obtain our final meta-analytic network. We then applied hierarchical clustering with average linkage and extracted modules using the dynamic cutting tree algorithm (Langfelder et al., 2008).

To compute the association between co-expression modules and projection patterns, we framed the association as a classification task: can we predict projection patterns from module expression? First, we generated labels by binarizing each projection pattern: cells with a projection strictly greater than the median projection strength were marked as positives. Then we generated predictors by computing gene module expression as the average Log(CPM+1) across all genes in the module. We reported the association strength (classification results) as an area under the receiver-operator characteristic curve (AUROC). To compute the association between co-expression modules and cell types, we used a similar approach, using clusters defined by the BICCN (Yao et al., 2020) as labels. For visualization, cell types are organized according to the following procedure: cell types are reduced to a centroid by taking the median expression for each gene, then cell types are clustered according to hierarchical clustering with average linkage with correlation-based distance.

### Statistical tests

All statistical tests performed were indicated in the main text. Bonferroni correction was used for all p values reported unless noted otherwise. Wherever indicated, False Discovery Rates (FDRs) were computed according to the Benjamini-Hochberg procedure (Benjamini and Hochberg, 1995).

### Data availability

Target area sequencing data are deposited at SRA (SRR12247894, SRR12245390, and SRR12245389). Raw *in situ* sequencing images are deposited at Brain Image Library. Other data and processing scripts are deposited at Mendeley Data (preview link: https://data.mendeley.com/datasets/jnx89bmv4s/draft?a=4672e3ab-4097-4a1d-8e1e-3729b5b5e3b6).

## Acknowledgement

The authors would like to acknowledge members of the MAPseq core facility, Huiqing Zhan, Yan Li, and Nicole Gemmill, for MAPseq data production, Katherine Matho and Z. Josh Huang for dissection coordinates in motor cortex, Huiqing Zhan, Li Yuan, Henry Lee Gilbert, Katherine Matho, Justus Kebschull, and Daniel Fürth for useful discussions, and Wiktor Wadolowski, Barry Burbach, Kathleen Lucere, and Eugene Fong for technical support. This work was supported by the National Institutes of Health [NIH 5RO1NS073129, 5RO1DA036913, RF1MH114132, and U01MH109113 to A.M.Z, R01MH113005 and R01LM012736 to J.G., and U19MH114821 to both A.M.Z. and J.G.], the Brain Research Foundation (BRF-SIA-2014-03 to A.M.Z.), IARPA MICrONS [D16PC0008 to A.M.Z.], Paul Allen Distinguished Investigator Award [to A.M.Z.], Simons Foundation [350789 to X.C.], Chan Zuckerberg Initiative (2017-0530 ZADOR/ALLEN INST(SVCF) SUB awarded to A.M.Z], and Robert Lourie (to A.M.Z.). This work was additionally supported by the Assistant Secretary of Defense for Health Affairs endorsed by the Department of Defense, 1120 Fort Detrick, Fort Detrick, MD 21702 through the FY18 PRMRP Discovery Award Program W81XWH1910083 awarded to X.C. Opinions, interpretations, conclusions and recommendations are those of the author and are not necessarily endorsed by the U.S. Army. In conducting research using animals, the investigator adheres to the laws of the United States and regulations of the Department of Agriculture.

## Author Contributions

Y.-C.S., X.C., and A.M.Z. conceived the study. Y.-C.S. and X.C. optimized and performed BARseq2. X.C., S.F. and J.G. analyzed data. Y.-C.S., X.C. and S.F. selected gene panels. X.C. and S. L. compared gene expression between BARseq2 and Allen ISH. Y.-C.S. and X.C. performed retrograde tracing combined with FISH validations. Y.-C.S., X.C., S.F., and A.M.Z. wrote the paper.

## Declaration of Interests

A.M.Z. is a founder and equity owner of Cajal Neuroscience and a member of its scientific advisory board.

## Notes

https://data.mendeley.com/datasets/jnx89bmv4s/draft?a=4672e3ab-4097-4a1d-8e1e-3729b5b5e3b6

