## Supplementary Note for "Integrating barcoded neuroanatomy with spatial transcriptional profiling reveals cadherin correlates of projections shared across the cortex"

**Supplementary Notes**

**Supplementary Note 1: Optimization of endogenous mRNA detection**

We optimized padlock probes, tissue pretreatment, and reverse transcription to maximize detection sensitivity. We found that using multiple padlocks per mRNA transcript, each padlock targeting a different site on the mRNA coding sequence, increased detection efficiency significantly (Fig. 1E). The increase in sensitivity varied between genes, but this is likely caused by differences in sensitivity of the single probe to which we normalized the sensitivities. For tissue pretreatment, we found that thin fresh frozen tissue cryosections fixed with 4% PFA for 30 mins to 1 hour (Extended Data Fig. 1A) yielded higher mRNA sensitivity than shorter fixation or other pretreatments, such as PFA-perfused tissue slices with or without post-fixation. For reverse transcription, we found that reverse transcription primers specific to the targets at a concentration of 0.5 – 5 µM each yielded higher sensitivity than using random primers at concentrations up to 50 µM (Extended Data Fig. 1B). Altogether, these optimizations were crucial for increased mRNA detection sensitivity comparable to hybridization-based techniques.

To quantify the sensitivity of BARseq2 compared to conventional FISH methods, we detected two genes, *Slc30a3* and *Cdh13*, using both BARseq2 and RNAscope (Fig. 1F). We also probed for a third gene, *Slc17a7*, but at the resolution we imaged at, we were unable to fully resolve the signals from both BARseq2 and RNAscope. We therefore only used *Slc30a3* and *Cdh13*, not *Slc17a7*, to evaluate the sensitivity of BARseq2. Linear regression between BARseq2 and RNAscope counts of *Slc30a3* and *Cdh13* genes in these two genes resulted in a slope of 1.65 (Extended Data Fig. 1C, D; dashed line R^2^ = 0.73), indicating that BARseq2 achieved about 1 / 1.65 ≈ 60% sensitivity compared to RNAscope.

**Supplementary Note 2: Laminar distribution of cadherins**

Because many genes, especially cell adhesion molecules, are differentially expressed across cortical layers, we evaluated how well BARseq2 can capture spatial organization of cadherins compared to existing methods, such as FISH.

RNAscope against *Cdh8*, *Pcdh19*, and *Pcdh20* revealed laminar expression profiles that were qualitatively similar to those obtained by BARseq2 (Fig. 2E). For *Pcdh20*, the dynamic range of gene expression (i.e. the differences between peaks and valleys in expression) was more pronounced in the BARseq2 data than that observed by RNAscope. Because low sensitivity and/or low specificity would likely result in a reduction, not an increase, in the dynamic range of expression, it is unlikely that such quantitative differences in the laminar profiles of gene expression were caused by sensitivity and/or specificity issues with BARseq2. We suspect that the reduced dynamic range in RNAscope is caused by non-specific signals inherent to amplified FISH methods. We therefore sought to compare BARseq2 to other FISH datasets to confirm its accuracy.

We then compared the distributions of genes obtained by BARseq2 to those in the Allen gene expression atlas (Lein et al., 2007)(Fig. 2F; Extended Data Fig. 3). The laminar distribution of gene expression revealed by BARseq2 was highly correlated with that in the Allen gene expression atlas (Spearman correlation ρ = 0.696, p = 3.8×10^-29^). Specifically, the laminar distribution of *Pcdh20* obtained by BARseq2 matched very well with *Pcdh20* in the Allen gene expression atlas (Extended Data Fig. 3). These results indicate that BARseq2 accurately captured the laminar distribution of cadherin expression.

**Supplementary Note 3: *Slc17a7* and *Gad1* expression observed by BARseq2**

*Slc17a7* and *Gad1* are expressed in excitatory and inhibitory neurons, respectively. They are thus almost never expressed in the same neuron in the cortex. BARseq2 recapitulated this mutual exclusivity between these two genes (Fig. 2J), but a small number of neurons did express both *Slc17a7* and *Gad1* (grey cells in Fig. 2J). This could be caused by overlapping cells (i.e. an inhibitory neuron and an excitatory neuron at the same x/y position, but in different z planes were merged together in the max projection images) or cell segmentation errors (two adjacent cells incorrectly segmented as a single cell). Because the sections we used were 10 µm thick, which was comparable to the diameter of an average neuron, the latter source of error was likely to be more common.

This type of error was similar to doublets in droplet-based single-cell RNAseq techniques. Assuming that the mutual exclusions of *Slc17a7* and *Gad1* were absolute, then we could estimate the “doublet” rate as the ratio between the probability of neurons expressing both genes and the product of the probabilities of neurons expressing either gene. Using this formula, we estimated the doublet rate of BARseq2 to be 7.5%, which is in a similar range as droplet-based single-cell RNAseq techniques (usually < 5%). Further improvement in cell segmentation algorithms may further reduce the doublet rate.

In addition to cells that express both *Gad1* and *Slc17a7* at significant levels, most cells that expressed one of the two genes dominantly also had non-zero expression of the other gene, albeit at much lower levels. This noise floor could be caused by mRNAs in dendrites that were incorrectly assigned to other neurons. Because the expression of these genes in the somata were much higher than that in the dendrites, this type of error was unlikely to significantly affect the determination of excitatory and inhibitory neurons.

**Supplementary Note 4: Cell type calling using a marker gene panel in M1**

BARseq2, like most other imaging-based spatial transcriptomic methods, relies on a selected panel of genes to determine transcriptomic cell types. To generate the panel of marker genes, we chose up to 5 genes for each transcriptomic type of excitatory neurons based on single-cell RNAseq in the motor cortex (Yao et al., 2020). These genes were selected based on first, the number of reference datasets in which the gene was detected as differentially expressed (fold change > 2, FDR < 0.05), and second, the average fold change across datasets. We additionally required genes to have a minimal expression level (average CPM > 100). Each cell was then assigned to the cell type with the maximum mean expression of marker genes compared to other cell types. This method of cell typing achieved good precision and recall for most cell types when applied to single-cell RNAseq data (Extended Data Fig. 5D). We applied the procedure across 9 datasets to check whether it is robust across technologies and sequencing depth (Extended Data Fig. 5E, F). Overall, we observed extremely high performance for NP and CT subtypes in all cases, while L6b was slightly better predicted in high depth datasets. The cell typing method always predicted IT cells correctly, but not always the correct layer (L2/3, L5, L6, Car3, Extended Data Fig. 5G). This is consistent with the observation that IT types form a continuum in single cell datasets, making it difficult to fully separate subtypes by layer. Finally, the PT type proved to be the most difficult type to predict. While all PT cells were correctly annotated as PT (Extended Data Fig. 5H), numerous L2/3 IT and L5 IT cells were wrongly annotated as PT, in particular in high depth datasets (Extended Data Fig. 5F, G). We believe that this was due to an imbalance in the marker panel, with PT markers being higher expressing than markers from other types. We tested various normalization procedure to overcome this effect but found that results were insensitive to normalization overall (Extended Data Fig. 5F).

Using this panel and cell typing method, we determined the transcriptomic types of excitatory neurons in motor cortex using BARseq2 (Fig. 3B). Most transcriptomic types were found enriched in the correct layers. One exception to this was the L6 Car3+ IT type. In general, very few L6 Car3+ IT neurons were identified by BARseq2. Furthermore, even though L6 Car3+ IT neurons were predominantly in L6, some were identified in L2/3 by BARseq2 (Fig. 3C). This result was surprising, given that L6 Car3+ IT neurons, when present, were only rarely mistyped as L2/3 in our preliminary analyses (Extended Data Fig. 5G). L6 Car3+ IT neurons were only rarely detected in the datasets used to select markers, so we expect that using additional data will lead to a more robust marker selection and better cell-typing performance with BARseq2. These optimizations, however, are beyond the scope of this paper.

**Supplementary Note 5: Excitatory and inhibitory projection neurons**

Consistent with previous observations, most cortical projection neurons identified by BARseq2 were excitatory. However, we also identified a small fraction of inhibitory projection neurons (Fig. 4E). Some of these neurons could be caused by “doublets” as discussed in Supplementary Note 3. Consistent with this hypothesis, the inhibitory projection neurons (and some excitatory projection neurons) in motor cortex expressed both *Gad1* and *Slc17a7* at similar levels (Extended Data Fig. 6G). However, inhibitory projection neurons in auditory cortex expressed only *Gad1*, not *Slc17a7* (Extended Data Fig. 6H), suggesting that these were real inhibitory projection neurons. This observation was consistent with previous reports of rare inhibitory projection neurons in the cortex (Chen et al., 2019; Rock et al., 2016). We did not further analyze these inhibitory projection neurons.

We also observed many excitatory neurons without projections (Fig. 4D, E), similar to those observed in previous BARseq experiments (Chen et al., 2019). These neurons were likely non-projecting excitatory neurons and neurons that project only locally or to neighboring cortical areas (Tasic et al., 2018) that we did not sample.

**Supplementary Note 6: Differential cadherin expression across major classes compared to single-cell RNAseq**

Major classes of projection neurons (IT, PT, and CT) differ in both gene expression and projection patterns. Therefore, the differential expression of cadherins observed across these three major classes defined by projection patterns should be consistent with the differential expression across the classes defined by transcriptomic methods. To test this, we compared the differences in mean expression of cadherins in the three classes in motor and auditory cortex observed by BARseq2 to those observed using single-cell RNAseq in neighboring cortical areas (V1 and ALM) (Tasic et al., 2018). Generally, differentially expressed cadherins identified by BARseq were also differentially expressed in single-cell RNAseq (Extended Data Fig. 7A). Importantly, all cadherins that were consistently differentially expressed in both A1 and M1 were also differentially expressed across the same pairs of major classes in V1 and ALM as shown by single-cell RNAseq (purple dots in Extended Data Fig. 7A). Several cadherins, including *Pcdh7* and *Cdh11*, were differentially expressed with the opposite signs in single-cell RNAseq and in BARseq2 (yellow dots in lower right quadrant in Extended Data Fig. 7A). However, these cadherins were not consistently expressed across motor and auditory cortex. For example, *Pcdh7* was expressed at significantly higher level in PT neurons than CT neurons in motor cortex (p < 10^-8^; Fig. 5B), but at lower level in PT neurons than CT neurons in auditory cortex (p = 0.0011, not statistically significant at FDR < 0.05). It is thus likely that these differences between observations by BARseq2 and by single-cell RNAseq reflect area-to-area differences, not methodological differences. These results confirm the differential expression of cadherins across major classes identified by BARseq2.

**Supplementary Note 7: Validation of cadherin correlates of IT projections using *in situ* hybridization and retrograde labeling.**

To confirm that *Cdh8*, *Cdh12*, and *Pcdh19* correlated with ipsilateral, contralateral, and striatal projections, respectively, we performed CTB retrograde labeling from the projection targets and performed FISH against *Slc17a7*, *Slc30a3*, and the cadherins in both A1 and M1 (Extended Data Fig. 8A). We then quantified cadherin expression and CTB labeling in IT neurons that had good DAPI signals and expressed both *Slc17a7*, an excitatory cell marker, and *Slc30a3*, which labeled the majority of IT neurons (Extended Data Fig. 8B). Neurons that had weak and/or ambiguous CTB signals were excluded from the analyses. Indeed, we saw that the three cadherins were expressed at higher levels in CTB+ neurons in both areas despite significant overlap in expression between CTB+ and CTB- neurons (Extended Data Fig. 8C-E). This overlap was expected because CTB was unlikely to have labeled all neurons that projected to the areas that we sampled with BARseq2. For example, in a previous study, we found that less than half of neurons with projections detected by BARseq were also labeled by CTB injected to the same target area (Chen et al., 2019). These results thus provide further support for the finding that cadherins correlate with similar projections in both A1 and M1.

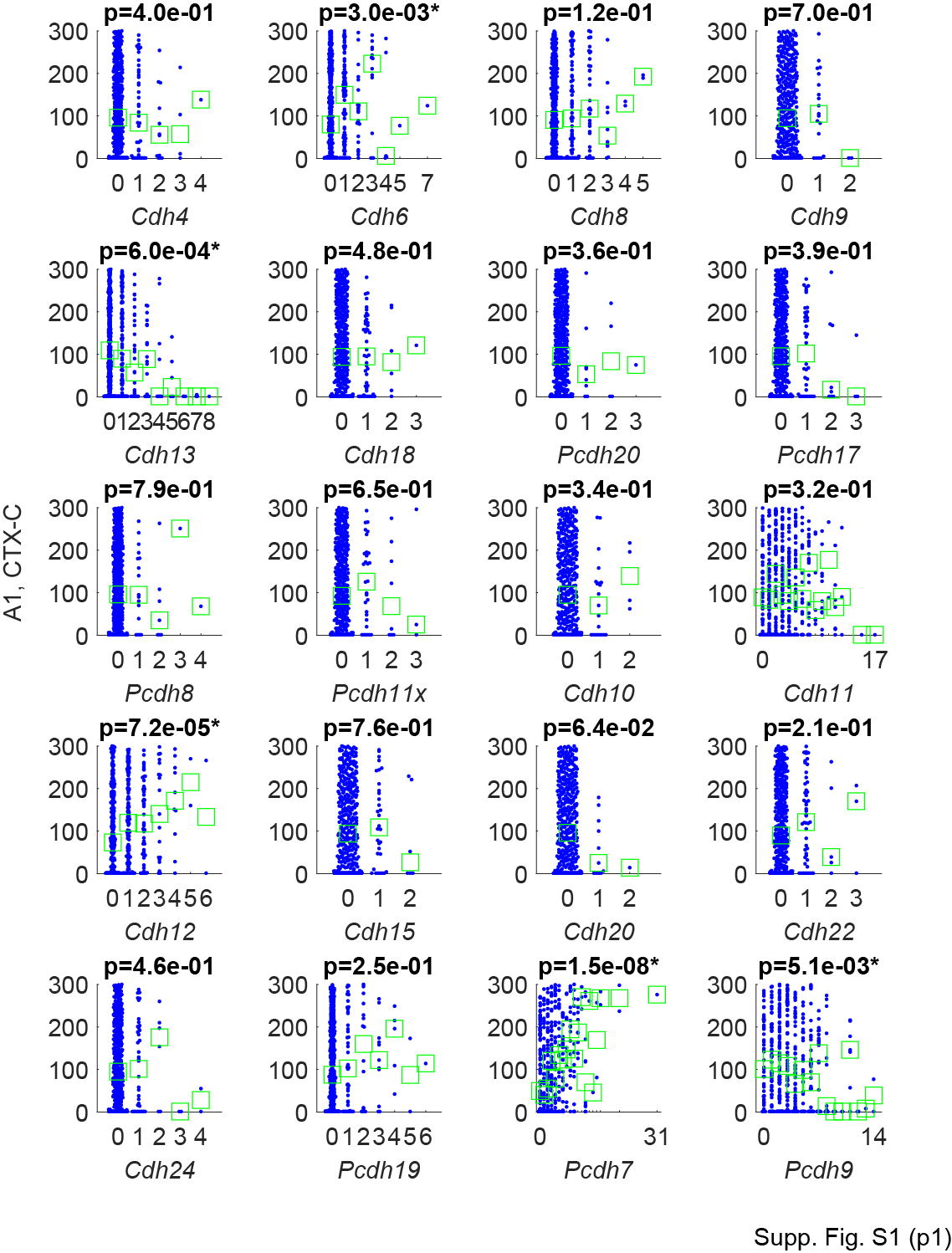

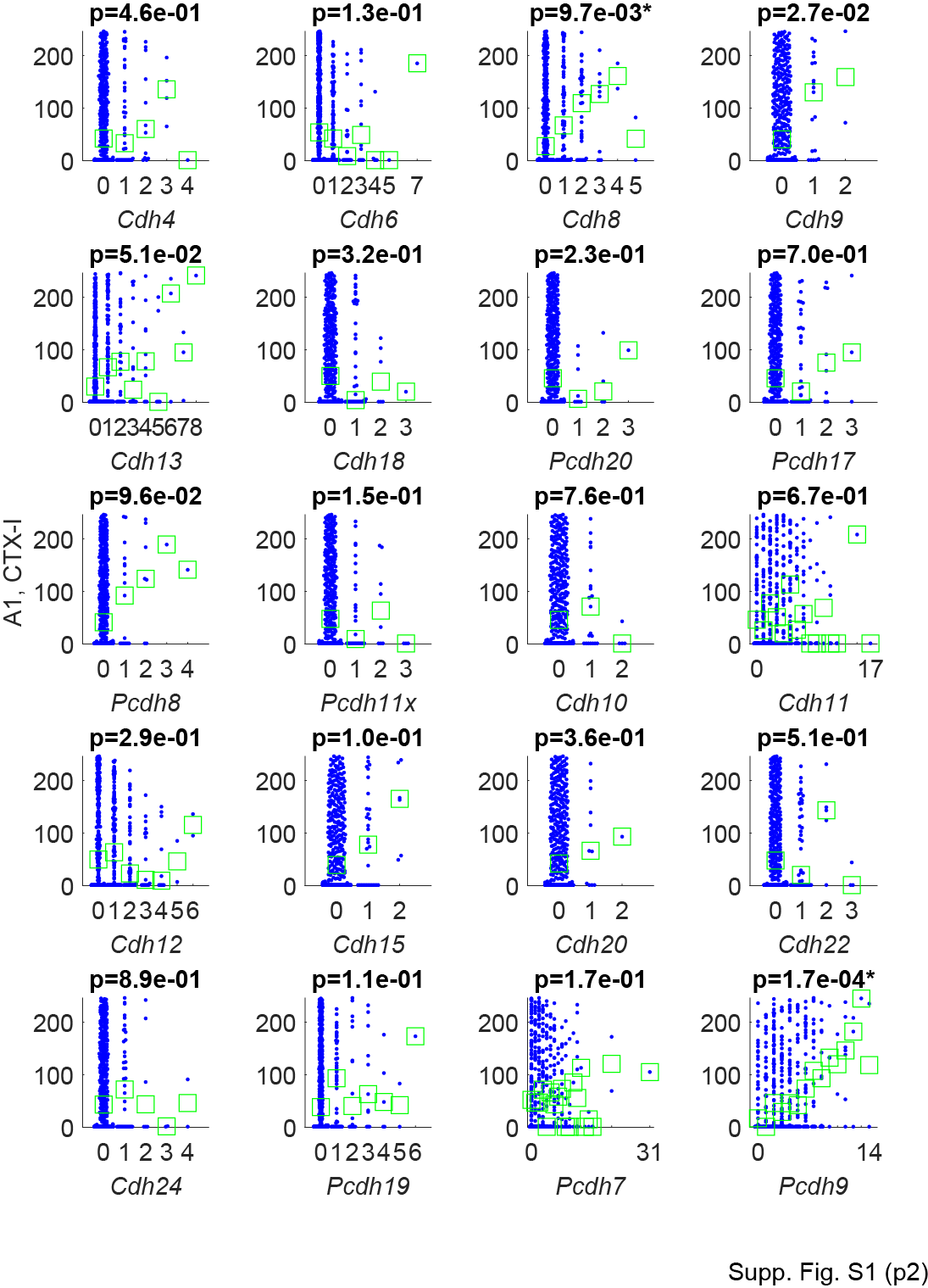

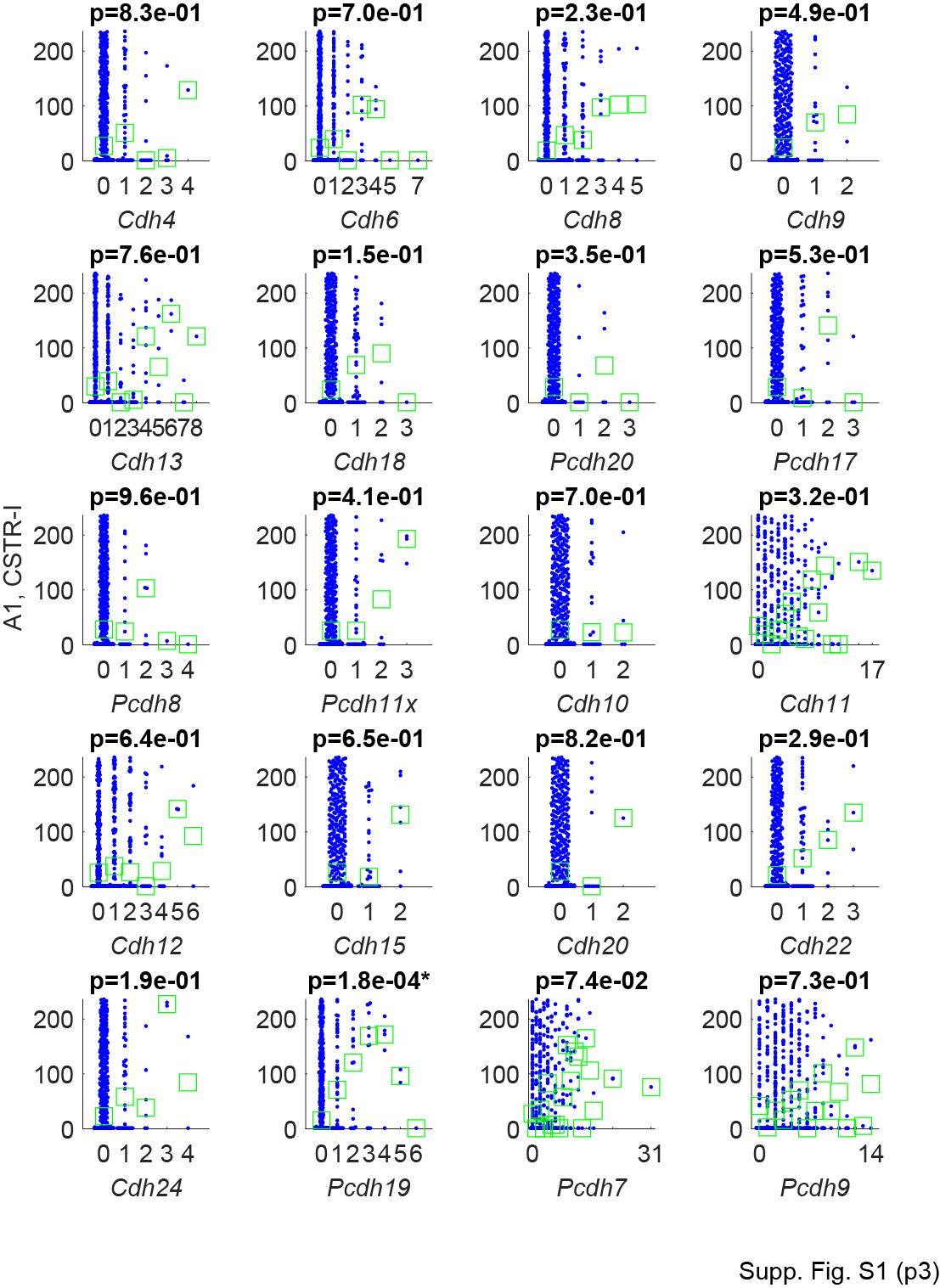

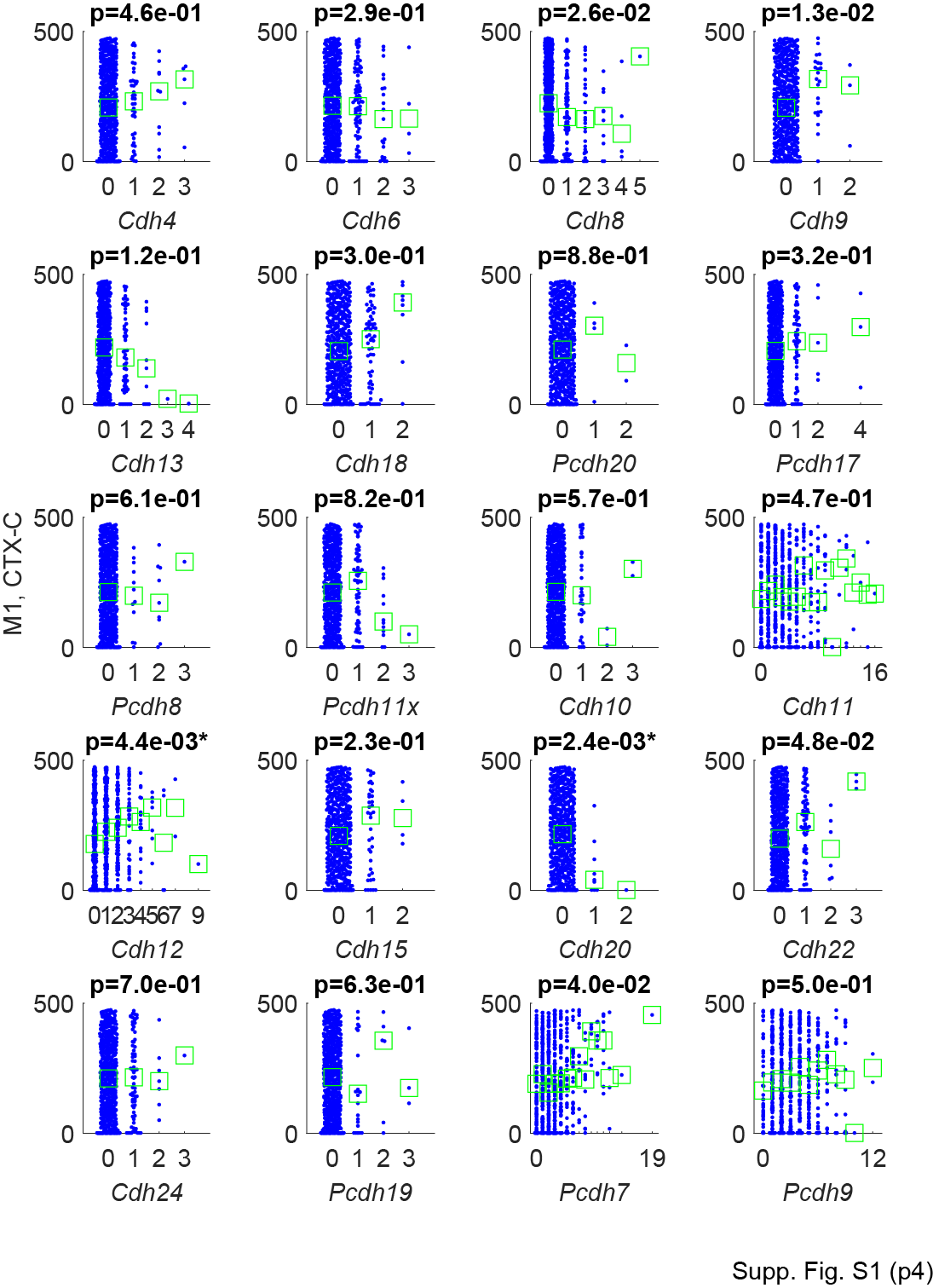

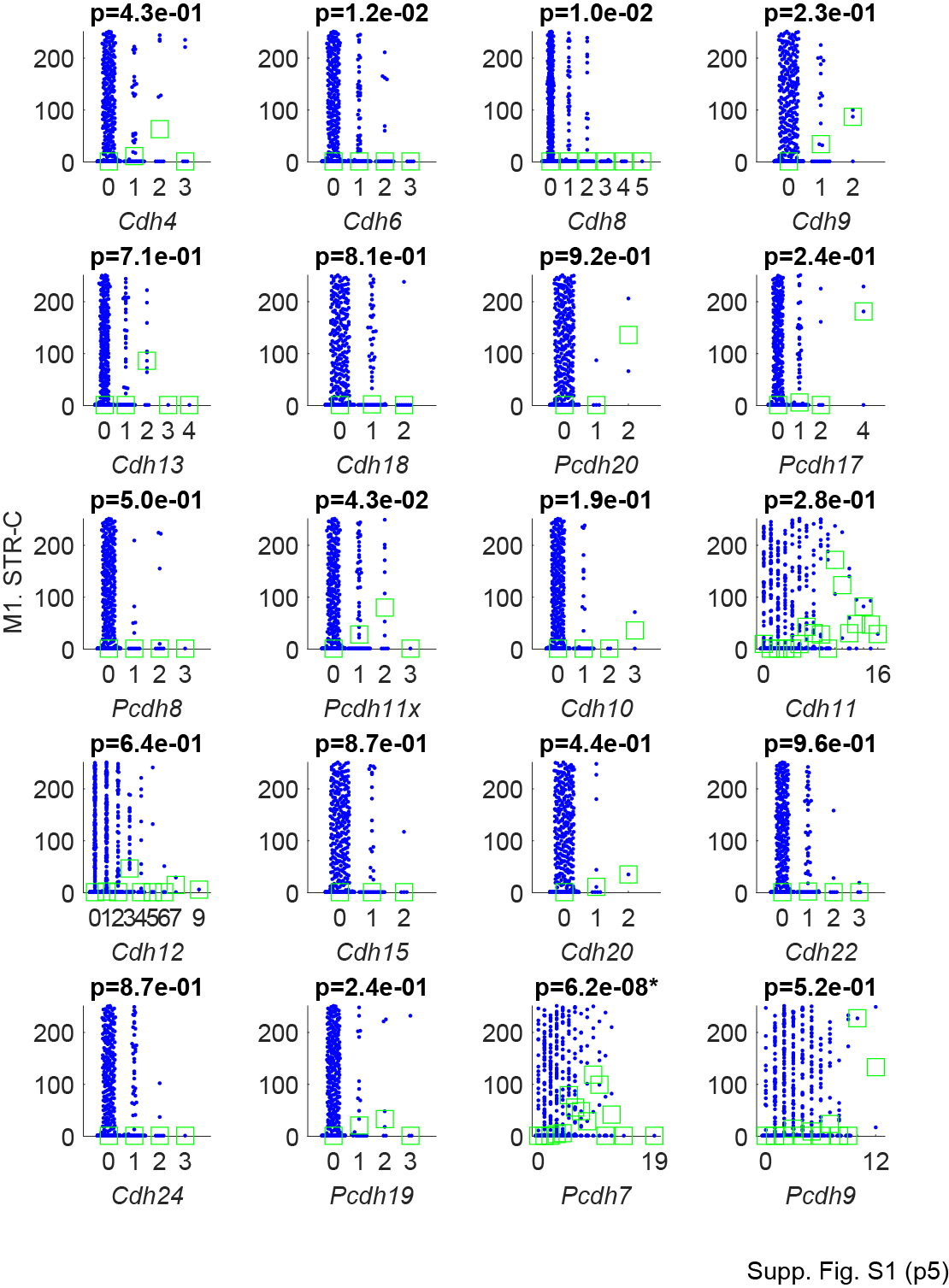

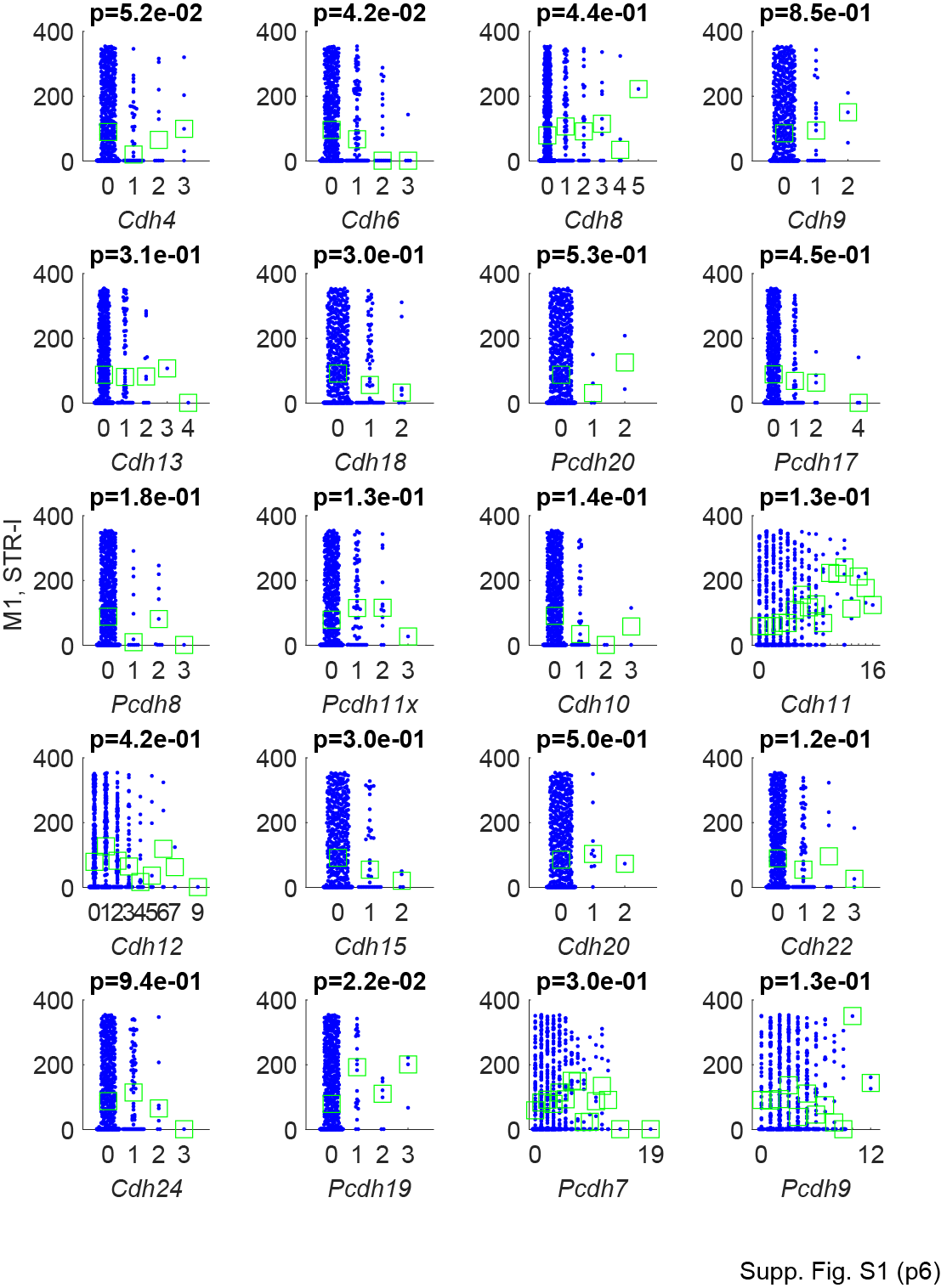

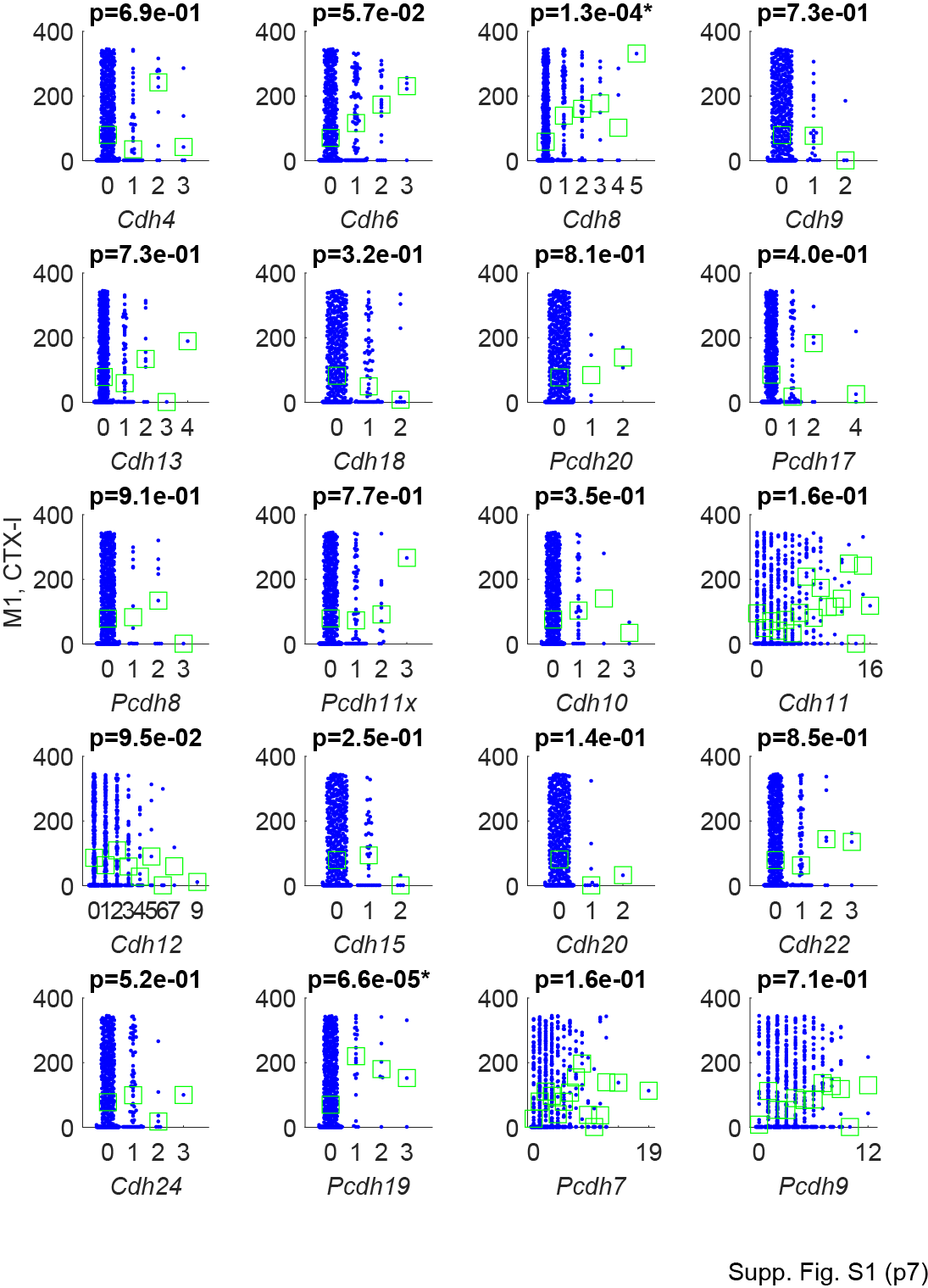

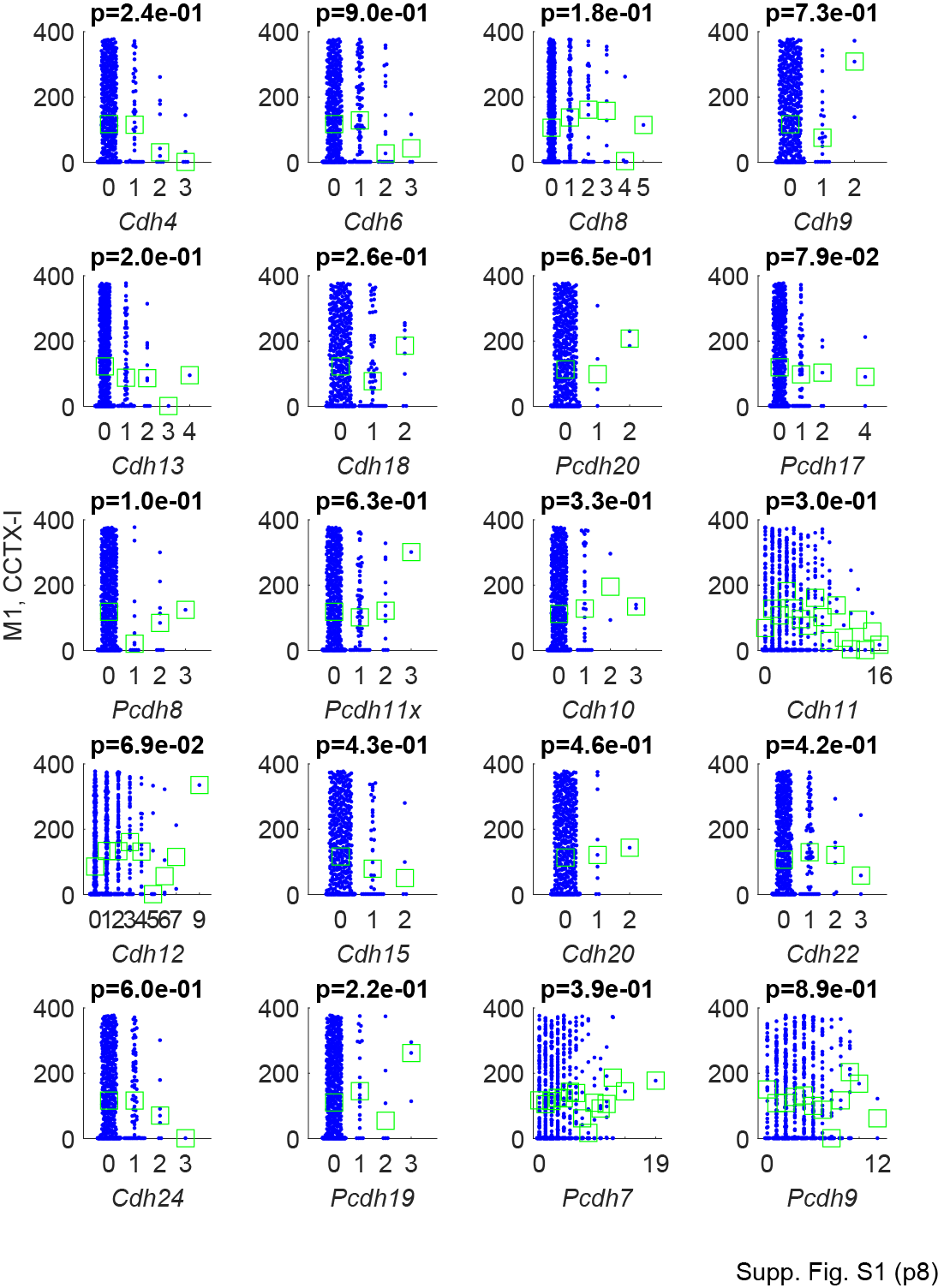

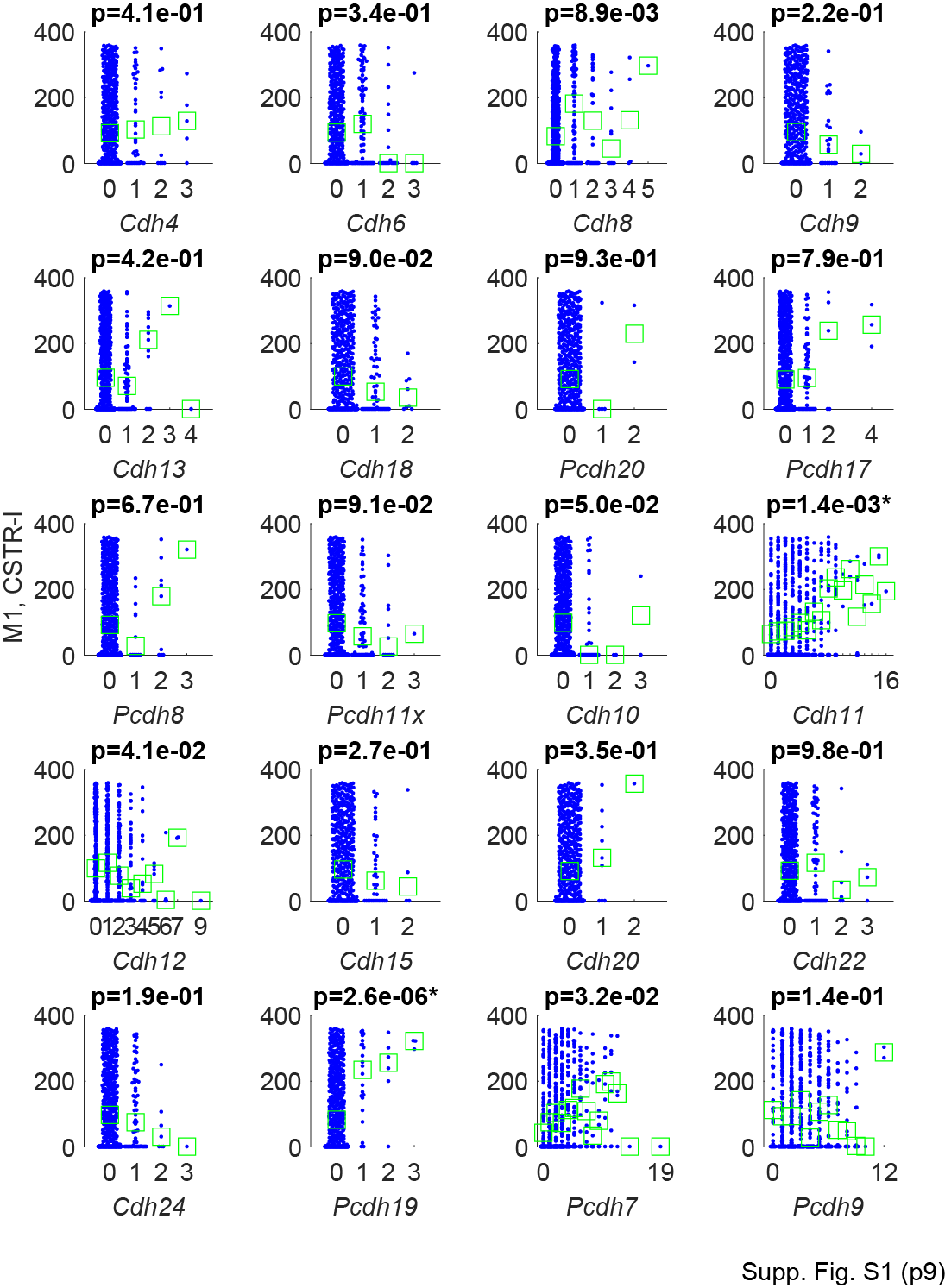

**Supplementary Figure S1.** The rank of the coefficients of the indicated projection modules (y-axes) for neurons with the indicated expression of cadherins (x-axes) in auditory cortex and motor cortex. Each dot represents a neuron. Median values are indicated by green squares. Raw p-values from rank sum tests are shown on top of each graph and those with FDR < 0.1 are labeled with an asterisk.

**(See SuppTableS1.xlsx)**

**Supplementary Table S1. The sequences of primers, probes, and GII codes used in the study.**

| **A1** | **BARseq area name** | **Allen area labels** | **Coronal level** | **Ipsi/contra** |
| --- | --- | --- | --- | --- |
|  | OB | MOB | / | Ipsilateral |
|  | OFC | ORBvl/ORBl/ORBm | 24-38 | Ipsilateral |
|  | M | MOs/MOp | 27-45 | Ipsilateral |
|  | RStr | CP | 42-56 | Ipsilateral |
|  | SS | SSp/SSs | 59-70 | Ipsilateral |
|  | CStr | CP | 64-70 | Ipsilateral |
|  | Amyg | COAa/COApl/PAA /IA/BMAa/BMAp | 64-70 | Ipsilateral |
|  | AudC | AUDd/AUDp/AUDv | 73-87 | Contralateral |
|  | VisC | PTLp | 73-87 | Contralateral |
|  | VisIp | PTLp | 73-87 | Ipsilateral |
|  | Thal | TH | 73-87 | Ipsilateral |
|  | Tect | SCs/SCm/IC | 87-116 | Ipsilateral |
| **M1** | **BARseq area name** | **Allen area labels** | **Coronal level** | **Ipsi/contra** |
|  | OB | MOB | / |  |
| Cortex | | | | |
|  | ORB-i | ORBvl/ORBl/ORBm | 31-37 | Ipsilateral |
|  | ORB-c | ORBvl/ORBl/ORBm | 31-37 | Contralateral |
|  | MOs-i | MOs | 23-35 | Ipsilateral |
|  | MOs-c | MOs | 23-35 | Contralateral |
|  | CLA-i | CLA | 40-46 | Ipsilateral |
|  | CLA-c | CLA | 40-46 | Contralateral |
|  | MOp-c | MOp | 44-59 | Contralateral |
|  | SS-i | SSs | 58-70 | Ipsilateral |
|  | SS-c | SSs | 58-70 | Contralateral |
|  | SSp-i | SSp | 60-69 | Ipsilateral |
|  | SSp-c | SSp | 60-69 | Contralateral |
|  | TEa-i | TEa/ECT/PERI/ENTl | 76-95 | Ipsilateral |
|  | TEa-c | TEa/ECT/PERI/ENTl | 76-95 | Contralateral |
| Striatum | | | | |
|  | Str-r-i | CP | 39-47 | Ipsilateral |
|  | Str-r-c | CP | 39-47 | Contralateral |
|  | Str-i-i | CP | 48-56 | Ipsilateral |
|  | Str-i-c | CP | 48-56 | Contralateral |
|  | Str-c-i | CP | 57-72 | Ipsilateral |
|  | Str-c-c | CP | 57-72 | Contralateral |
| Thalamus | | | | |
|  | Thal-mr-i | TH | 63-69 | Ipsilateral (medial half) |
|  | Thal-lr-i | TH | 63-69 | Ipsilateral (lateral half) |
|  | Thal-mr-c | TH | 63-69 | Contralateral (medial half) |
|  | Thal-lr-c | TH | 63-69 | Contralateral (lateral half) |
|  | Thal-mc-i | TH | 73-80 | Ipsilateral (medial half) |
|  | Thal-lc-i | TH | 73-80 | Ipsilateral (lateral half) |
|  | Thal-mc-c | TH | 73-80 | Contralateral (medial half) |
|  | Thal-lc-c | TH | 73-80 | Contralateral (lateral half) |
|  | ZI-i | ZI | 80-83 | Ipsilateral |
| Midbrain | | | | |
|  | MRN | MRN/RN/PAG/SC | 83-89 | Ipsilateral |
|  | SC | SC | 96-103 | Ipsilateral |
|  | PG | PG | 90-97 | Ipsilateral |
| Medulla | | | | |
|  | MY-l-i | MY | 108-132 | Ipsilateral (lateral half) |
|  | MY-m-i | MY | 108-132 | Ipsilateral (medial half) |
|  | MY-l-c | MY | 108-132 | Contralateral (lateral half) |
|  | MY-m-c | MY | 108-132 | Contralateral (medial half) |
| Spinal cord | | | | |
|  | Sp | NA | NA | NA |

**Supplementary Table S2. List of area abbreviations with the corresponding coronal levels and area names in the Allen reference atlas.**

| Date | ID | Experiment | Injection | Strain | Processing | MAPseq |
| --- | --- | --- | --- | --- | --- | --- |
| 5/30/2019 | XC95 | A1 BARseq1 | -2.5 mm AP, -4.3 mm ML, 300 600 900 µm DV, 30degree, 180 nL 1:3 diluted | C57BL/6J, 8 wks, Charles River | 24 hrs post injection, fresh frozen for bulk NGS sequencing. Injection site punched out with 2mm punch and fresh frozen for in situ sequencing | Mseq063_ZL182 |
| 8/19/2019 | XC97 | A1 BARseq2 | -2.5 mm AP, -4.3 mm ML, 300 600 900 µm DV, 30degree, 180 nL 1:3 diluted | C57BL/6J, 8 wks, Charles River | 24 hrs post injection, fresh frozen for bulk NGS sequencing. Injection site punched out with 2mm punch and fresh frozen for in situ sequencing | Mseq079_ZL189 |
| 11/19/2019 | XC116 | M1 BARseq | 0.5 mm AP, -1.5 mm ML, 500 900um DV, 180 nL 1:3 diluted | C57BL/6J, 9 wks, Charles River | 24 hrs post injection, fresh frozen for bulk NGS sequencing. Injection site punched out with 2mm punch and fresh frozen for in situ sequencing | Mseq102_ZL207 |
| 1/24/2020 | EF146B | A1 contralateral CTB | -2.88 mm AP, -4.3 mm ML, 300 600 900 µm DV, 130 nL 1.0 mg/mL | C57BL/6J, 8 wks, Jackson Laboratory | 96 hrs post injection, PFA perfusion, 24 hrs postfix, cryoprotection. | NA |
| 6/11/2020 | XC160 | A1 contralateral CTB | -2.88 mm AP, -4.3 mm ML, 300 600 900 µm DV, 130 nL 1.0 mg/mL | C57BL/6J, 8 wks, Jackson Laboratory | 96 hrs post injection, PFA perfusion, 24 hrs postfix, cryoprotection. | NA |
| 6/12/2020 | XC162 | A1 ipsilateral CTB | -2.555 mm AP, -1.0 mm ML, 300 µm DV, 90 nL 1.0 mg/mL | C57BL/6J, 8 wks, Jackson Laboratory | 96 hrs post injection, PFA perfusion, 24 hrs postfix, cryoprotection. | NA |
| 6/9/2020 | XC166 | A1 ipsilateral CTB | -2.555 mm AP, -1.0 mm ML, 300 µm DV, 90 nL 1.0 mg/mL | C57BL/6J, 8 wks, Jackson Laboratory | 96 hrs post injection, PFA perfusion, 24 hrs postfix, cryoprotection. | NA |
| 6/9/2020 | XC168 | A1 striatum CTB | -1.555 mm AP, -3.5 mm ML, 3.75 mm DV, 90 nL 1.0 mg/mL | C57BL/6J, 8 wks, Jackson Laboratory | 96 hrs post injection, PFA perfusion, 24 hrs postfix, cryoprotection. | NA |
| 7/3/2020 | XC179 | A1 striatum CTB | -1.555 mm AP, -3.5 mm ML, 3.5 mm DV, 90 nL 1.0 mg/mL | C57BL/6J, 8 wks, Jackson Laboratory | 96 hrs post injection, PFA perfusion, 24 hrs postfix, cryoprotection. | NA |
| 3/12/2020 | YSMo1 | M1 contralateral CTB | 0.445 mm AP, -1.75 mm ML, 300 600 900 µm DV, 130 nL 1.0 mg/mL | C57BL/6J, 8 wks, Jackson Laboratory | 96 hrs post injection, PFA perfusion, 24 hrs postfix, cryoprotection. | NA |
| 3/12/2020 | YSMo2 | M1 contralateral CTB | 0.445 mm AP, -1.75 mm ML, 300 600 900 µm DV, 130 nL 1.0 mg/mL | C57BL/6J, 8 wks, Jackson Laboratory | 96 hrs post injection, PFA perfusion, 24 hrs postfix, cryoprotection. | NA |
| 5/20/2020 | YSMoIpsi | M1 ipsilateral CTB | 2.445 mm AP, -0.75 mm ML, 300 600 µm DV, 90 nL 1.0 mg/mL | C57BL/6J, 8 wks, Jackson Laboratory | 96 hrs post injection, PFA perfusion, 24 hrs postfix, cryoprotection. | NA |
| 6/11/2020 | XC158 | M1 ipsilateral CTB | 2.445 mm AP, -0.75 mm ML, 300 600 µm DV, 90 nL 1.0 mg/mL | C57BL/6J, 8 wks, Jackson Laboratory | 96 hrs post injection, PFA perfusion, 24 hrs postfix, cryoprotection. | NA |
| 6/8/2020 | XC163 | M1 striatum CTB | -1.155 mm AP, -3.25 mm ML, 3.5 mm DV, 90 nL 1.0 mg/mL | C57BL/6J, 8 wks, Jackson Laboratory | 96 hrs post injection, PFA perfusion, 24 hrs postfix, cryoprotection. | NA |
| 6/8/2020 | XC164 | M1 striatum CTB | -1.155 mm AP, -3.25 mm ML, 3.5 mm DV, 90 nL 1.0 mg/mL | C57BL/6J, 8 wks, Jackson Laboratory | 96 hrs post injection, PFA perfusion, 24 hrs postfix, cryoprotection. | NA |

**Supplementary Table S3. Metadata of animals used in this study.**

| Microscope | Channel | Laser (nm) | Excitation | Dichroic | Emission (Filter or collection wavelength) |
| --- | --- | --- | --- | --- | --- |
| Spinning disk confocal | DAPI | 405 | zet402/468/555/640x (Chroma) | Zt402/468/555/640rpc-ufs (Chroma) | Zet402/648/555/640m (Chroma) |
| Spinning disk confocal | G/YFP | 520 | Zet443-518x (Chroma) | Zt443-518rpc (Chroma) | FF01-565/24 (Semrock) |
| Spinning disk confocal | T/RFP | 555 | zet402/468/555/640x (Chroma) | Zt402/468/555/640rpc-ufs (Chroma) | FF01-585/11 (Semrock) |
| Spinning disk confocal | A/Cy5 | 640 | zet402/468/555/640x (Chroma) | FF652-Di01 (Semrock) | FF01-676/29 (Semrock) |
| Spinning disk confocal | C | 640 | zet402/468/555/640x (Chroma) | FF652-Di01 (Semrock) | FF01-725/40 (Semrock) |
| Spinning disk confocal | GFP | 470 | zet402/468/555/640x (Chroma) | Zt402/468/555/640rpc-ufs (Chroma) | FF01-525/30 (Semrock) |
| Spinning disk confocal | TexasRed | 555 | zet402/468/555/640x (Chroma) | Di02-R594 (Semrock) | FF01-647/57 (Semrock) |
| LSM710 | GFP | 488 | N/A | MBS 488/561/633 | 494-562 nm |
| LSM710 | RFP | 561 | N/A | MBS 488/561/633 | 562-591 nm |
| LSM710 | TexasRed | 594 | N/A | MBS 488/594 | 596-740 nm |
| LSM710 | Cy5 | 633 | N/A | MBS 488/561/633 | 613-755 nm |
| LSM710 | DAPI | 405 | N/A | MBS-405 | 410-497 nm |

**Supplementary Table S4. Filter and laser settings used for imaging**
